# The development of attentional control mechanisms in multisensory environments

**DOI:** 10.1101/2020.06.23.166975

**Authors:** Nora Turoman, Ruxandra I. Tivadar, Chrysa Retsa, Anne M. Maillard, Gaia Scerif, Pawel J. Matusz

## Abstract

Outside the laboratory, people need to pay attention to relevant objects that are typically multisensory, but it remains poorly understood how the underlying neurocognitive mechanisms develop. We investigated when adult-like mechanisms controlling one’s attentional selection of visual and multisensory objects emerge across childhood. Five-, 7-, and 9-year-olds were compared with adults in their performance on a computer game-like multisensory spatial cueing task, while 129-channel EEG was simultaneously recorded. Markers of attentional control were behavioural spatial cueing effects and the N2pc ERP component (analysed traditionally and using a multivariate electrical neuroimaging framework). In behaviour, adult-like visual attentional control was present from age 7 onwards, whereas multisensory control was absent in all children groups. In EEG, multivariate analyses of the activity over the N2pc time-window revealed stable brain activity patterns in children. Adult-like visual-attentional control EEG patterns were present age 7 onwards, while multisensory control activity patterns were found in 9-year-olds (albeit behavioural measures showed no effects). By combining rigorous yet naturalistic paradigms with multivariate signal analyses, we demonstrated that visual attentional control seems to reach an adult-like state at ~7 years, before adult-like multisensory control, emerging at ~9 years. These results enrich our understanding of how attention in naturalistic settings develops.

**Highlights:**

- By age 7, children showed adult-like task-set contingent attentional capture in behaviour
- Children’s behavioural data did not show evidence for attentional enhancement for multisensory objects, but 9-year-olds’ EEG topographic patterns elicited by multisensory vs. purely visual distractors differed reliably
- Traditional visual attentional event-related potential (ERP) analyses, such as the N2pc, did not detect attentional enhancement for multisensory objects in adults, and visual or multisensory attention in children
- Multivariate analyses of ERPs, such as electrical neuroimaging, are more sensitive to the change of attentional control processes over development

## Introduction

Everyday environments contain many objects, so it is important to select only the relevant ones. Objects in such environments are also multisensory in nature. Here, we investigated whether adults and children pay attention to visual and multisensory stimuli in a similar way, and through similar brain mechanisms.

### 1. Everyday environments are multisensory

The brain integrates information across the senses, and so processes multisensory stimuli differently than unisensory stimuli. Multisensory processes lead to faster and more accurate behavioural responses (Stein & Meredith 1993; Murray & Wallace 2012) and improve learning and memory (Bahrick & Lickliter 2000; Lewkowicz, 2014). However, most studies focus on attentional control mechanisms engaged by unisensory (often visual) stimuli. This left aves unclear if attentional control mechanisms operate similarly on unisensory and multisensory stimuli. The relationships between multisensory integration and attentional control are a topic of ongoing debate (e.g. Talsma et al., 2010; van der Burg et al. 2011; Matusz et al. 2015, 2019a, 2019b). Previously, Matusz and Eimer (2011) has found that task-irrelevant multisensory distractors capture attention more strongly than visual distractors in an audiovisual adaptation of the Folk et al. (1992) spatial cuing task. In the Folk et al (1992) task, participants search for a target of a predefined colour (e.g., a red bar) within a multi-stimulus array. The array is always preceded by a distractor that either matches the target by colour (red) or a different, nontarget colour (blue). Colour distractors capture attention but only when they match the target colour – the so-called *task-set contingent attentional capture* hypothesis (Folk et al., 1992; see also Folk & Remington, 1998). Additionally, in In Matusz and Eimer (2011)’s study, on half of the trials the visual distractors were accompanied by a spatially-diffuse tone. Visual distractors captured attention more strongly when accompanied by sounds, both when they matched the target colour and when they did not (*multisensory enhancement of attention capture*). These findings suggest that findings from purely unisensory attentional research may be limited in explaining how attention to objects in space is controlled in real-world, multisensory settings. While we know relatively little about how adults control their attention towards multisensory objects, we know even less about how children do so, and if adults and children attend to multisensory objects using similar mechanisms.

### 2. Developing attentional control is poorly understood

Typically children have weaker visual attentional control skills than adults (Donnelly et al., 2007; Trick & Enns, 1998; Gaspelin et al., 2015). This can be due to the protracted development of the prefrontal and parietal cortices (Casey, et al., 2005; Tsujimoto, 2008), and the connectivity between them (e.g., Konrad et al., 2005; Hwang et al., 2010). However, the neurocognitive mechanisms underpinning these age-based differences are still unclear. For instance, children’s weak attentional control skills may be driven by weak interactions between top-down control and memory processes (e.g. Shimi et al. 2014a) or weak inhibitory mechanisms towards salient distractors (e.g. Hommel et al., 2004).

One way to create more complete and accurate theories of development of attention is by comparing the cognitive *and* brain mechanisms that children and adults use when paying attention to objects in naturalistic, multi-stimulus contexts (where location of the target (unlike other target attributes), is typically unknown. Such investigations are necessary for several reasons. First, in that context, one can benefit from using rigorous paradigms isolating relevant attentional control processes, such as the Folk et al. (1992)’s task. Second, the value of process-specific tasks is further increased when combined with well-researched and well-understood brain correlates, such as the N2pc event-related potential (ERP) component for attentional selection of objects in space (e.g. Eimer et al. 1996; Eimer et al. 2009). Indeed, adult N2pc studies have provided important corroborating evidence for task-set contingent attentional capture (e.g., Kiss et al., 2008; Eimer et al., 2009). An approach combining the two methods offers a useful ‘bridge’ in understanding the differences in how adults and children pay attention, as differences that may be more readily visible at the brain level (Astle & Scerif 2011). However, to date, the N2pc has been studied almost exclusively in adults. In rare exceptions, developmental N2pc studies revealed that the ability to focus on target-relevant objects among distractors is already adult-like in 10-year-olds and may rely on at least partly similar brain mechanisms to adults’ (e.g., visual search targets, Couperus & Quirk, 2015; targets held in memory, Shimi et al. 2015). In contrast, the development of other specific processes, like the task-set contingent attentional capture, has been little investigated (e.g., Gaspelin et al., 2015, in behaviour only). Thus, studying in children processes that are well understood in adults help map when children start using specific, adult-like mechanisms when paying attention. Third, as research in the area has been focused on studying alertness, orienting and executive attention processes separately (e.g., Rueda et al. 2004), paradigms that combine them in a rigorous fashion (e.g. spatial [attention capture] and featural [target colour] in Folk et al. 1992) should be particularly powerful in informing how attentional control develops. Finally, all of the above research focused on processes engaged by visual-only objects. Consequently, we have a limited understanding of neuro-cognitive mechanisms governing attention towards task-relevant and task-irrelevant multisensory objects, in children and adults alike (except one rare study on N2pc to audiovisual targets and distractors, van der Burg et al. 2011). Testing such mechanisms across adults and children can help build better theories of attentional control and how it develops.

### 3. Are children more sensitive to multisensory distraction?

Like all neurocognitive processes, multisensory processes undergo development. Brains are sensitive to congruence of stimulus onset, intensity, or identity already in infancy (Lewkowicz & Turkewitz, 1980; Bahrick & Lickliter, 2000; Neil et al. 2006). Other processes, related to perceptual judgements or sensorimotor skills mature slowly (8 and 10-11 years, Gori et al. 2008, 2012; and Barutchu et al. 2009, respectively). At the same time, the benefits of the multisensory nature of information for learning have been reported already at 5 years (Broadbent et al. 2018, 2019). Yet, existing research offers no insights as to whether children are more or less sensitive than adults to multisensory distraction. At what age do children and adults use similar neurocognitive mechanisms to attend to visual *and* multisensory information? As indirect evidence, we have previously shown that audiovisual distractors interfere with visual search in adults and 11-year-olds, but not 6-year-olds, for both colour (Matusz et al. 2015) and numerals (Matusz et al. 2019). In these studies, however, distractors were always task-relevant, as they shared the target’s identity. Thus, it remains to be established whether children are more or less sensitive than adults to completely task-irrelevant stimuli.

### 4. The present study

We developed a child-friendly version of the multisensory adaptation of Folk et al.’s spatial cueing task (Matusz & Eimer 2011), and tested it systematically on 5-, 7-, and 9-year-olds, as well as on young adults, while also recording their EEG. Through the use of colour-defined visual distractors and addition of sound on 50% of all trials as in the original Matusz and Eimer’s (2011) study, we could systematically investigate the differences between adults and children in controlling their attention towards visual and audiovisual objects, respectively. Specifically, we were interested in which of the two attentional control processes (visual or multisensory) reaches an adult-like state earlier.

We also investigated whether electrical neuroimaging (EN) analyses of the N2pc component are better at capturing developmental changes in attentional control over multisensory stimuli than the traditional N2pc analyses. Briefly, an EN approach encompasses a set of multivariate reference-independent analyses of the global features of the electric field measured at the scalp (Lehmann & Skrandies 1980; Murray et al., 2008; Tivadar & Murray 2019). The main added value of EN analyses is its ability to reveal the neurophysiological mechanisms driving differences in ERPs elicited between two (or more) experimental conditions, as arising due to differences in the strength of activation within a non-distinguishable brain network between conditions vs. differences in the networks recruited for responses between different conditions. Furthermore, classical N2pc analyses take into account much less EEG data than EN analyses, which suggests that EN analyses may detect effects that the canonical N2pc analyses may miss. In combination with rigorous paradigms and analyses of well-understood EEG correlates of cognitive processes, an EN approach offers a powerful tool to distinguishing between different accounts of cognitive processes, including multisensory attentional control (Matusz et al. 2019b; Turoman et al. 2020a).

We had the following predictions. Behaviourally, in adults, we expected to replicate task-set contingent attentional capture (“TAC”) and multisensory enhancement of attentional capture (“MSE”; see Matusz & Eimer, 2011). In children, we expected to find TAC at least in older groups (Gaspelin et al. 2015), without clear age-group predictions for MSE. For canonically measured N2pc, in adults, we expected to replicate TAC, indexed by attenuated/eliminated N2pc for non-target matching distractors. For MSE in N2pc, we did not have strong predictions, as the only related study to date showed little evidence for N2pc to audiovisual distractors (Van der Burg et al. 2011). In children, we did not have strong predictions for TAC and MSE effects as indexed by the N2pc. This is because, first, our oldest child group was younger than the youngest age group where N2pc was reported (9 years old here vs. 11 years old in e.g. Couperus & Quirk 2015). Second, the N2pc in our study was recorded in response to distractors, and not to targets, as in all the previous child studies. For EN analyses of the EEG within the N2pc time-window, we predicted that they will reveal modulations of brain responses by visual or multisensory attentional control in adults, and at least in older groups of children.

## Methods

### 1. Participants

A total of 39 young adults and 115 primary school children participated in the study. The children sample consisted of 28 children from the fifth grade, 46 children from the third grade, and 41 from the first grade of primary schools located in the canton of Vaud, Switzerland. In the local school system, children enter formal education at age 4, but only transition to sitting at desks instead of open seating and receiving less play-oriented instruction in third grade, when they are aged 6-7 years. To reduce confusion due to school system specificities, each child group is henceforth referred to by their members’ majority age, that is: ‘5-year-olds’ (first grade), ‘7-year-olds’ (third grade), and ‘9-year-olds’ (fifth grade). Children were recruited from local schools, nurseries, public events and entertainment facilities. Recruitment took place from March 2017 to May 2019. Of the total number of children recruited, first, 18 were excluded due to failure to initiate the testing session or failure to complete the task with above chance-level accuracy (50%), thus leading to exclusion of one 9-year-old, six 7-year-olds, and eleven 5-year-olds. Additionally, five additional participants (one 9-year-old, two 7-year-olds, and two 5-year-olds) were excluded because of unusable EEG signals due to excessive noise. Data for adult “controls” was taken from one task that was part of a larger study. The final sample consisted of 26 9-year-olds (10 male, *M_age_:* 8y 10mo, SD: 5mo, range: 8y 1mo – 10y 1mo), 38 7-year-olds (18 female, *M_age_:* 6y 10mo, SD: 4mo, range: 6y 1mo, 7y 9mo), and 28 5-year-olds (13 female, *M_age_:* 5y, SD: 4mo, range: 4y– 5y 7mo), and 39 adults (14 male, *M_age_:* 27y 6mo, SD: 4y, range: 22–38y).

Participants of all ages had normal or corrected-to-normal vision and normal hearing, and had no history of sensory problems (e.g., related to vision or audition), neurological problems (e.g., epilepsy), neurodevelopmental disorders (e.g., autism, ADHD), or learning difficulties (e.g., dyslexia), as indicated by parental report for children, or by direct report for adults. No children had an FSIQ under 70 which would warrant exclusion, as confirmed by an overall cognitive functioning assessment (see below). All research procedures were approved by the Cantonal Commission for the Ethics of Human Research (CER-VD). Informed consent was obtained from parents/caregivers and verbal assent was obtained from children before participating in the study.

### 2. Stimuli and procedure

All participants were tested at the Lausanne University Hospital Centre (CHUV). For children, the EEG session lasted between 1h and 1h30mins, including briefing, obtaining consent, the testing protocol, and breaks. For adults, the session took approximately 3h (part of a larger study), but the data used here were acquired within the first 1h-1h30, akin to the child protocol. Children’s baseline overall cognitive level was also assessed (although we do not report those results here), during a separate session on a different day. To this end, we used the Wechsler scale of intelligence for school-age (WISC-V, Wechsler, 2014) or preschool (WPPSI-IV, Wechsler, 2012) children, depending on the participant’s age. We used the abbreviated full-scale intellectual quotient (FSIQ) that included 4 subscales: Vocabulary, Matrix reasoning, Blocks and Similarities. After completing both sessions, children received a 30 Swiss franc voucher for a media store and parents/caregivers’ travel costs were reimbursed.

In the EEG session, participants completed a computer-game-like task where they searched for an elongated target diamond of a predefined colour (e.g. a blue diamond; see Figure 1), to help a pirate captain find treasure on a deserted island. Participants were instructed to assess the target diamond’s orientation (horizontal or vertical; randomly determined for each trial) and respond as quickly and accurately as possible by pressing one of two horizontally aligned round buttons (Lib Switch, Liberator Ltd.) that were fixed onto a tray bag on their lap. The search array was always preceded by an array containing a cue, which could match the target colour (e.g. blue set of dots) or be of another, nontarget colour (red set of dots).

**Figure 1.**
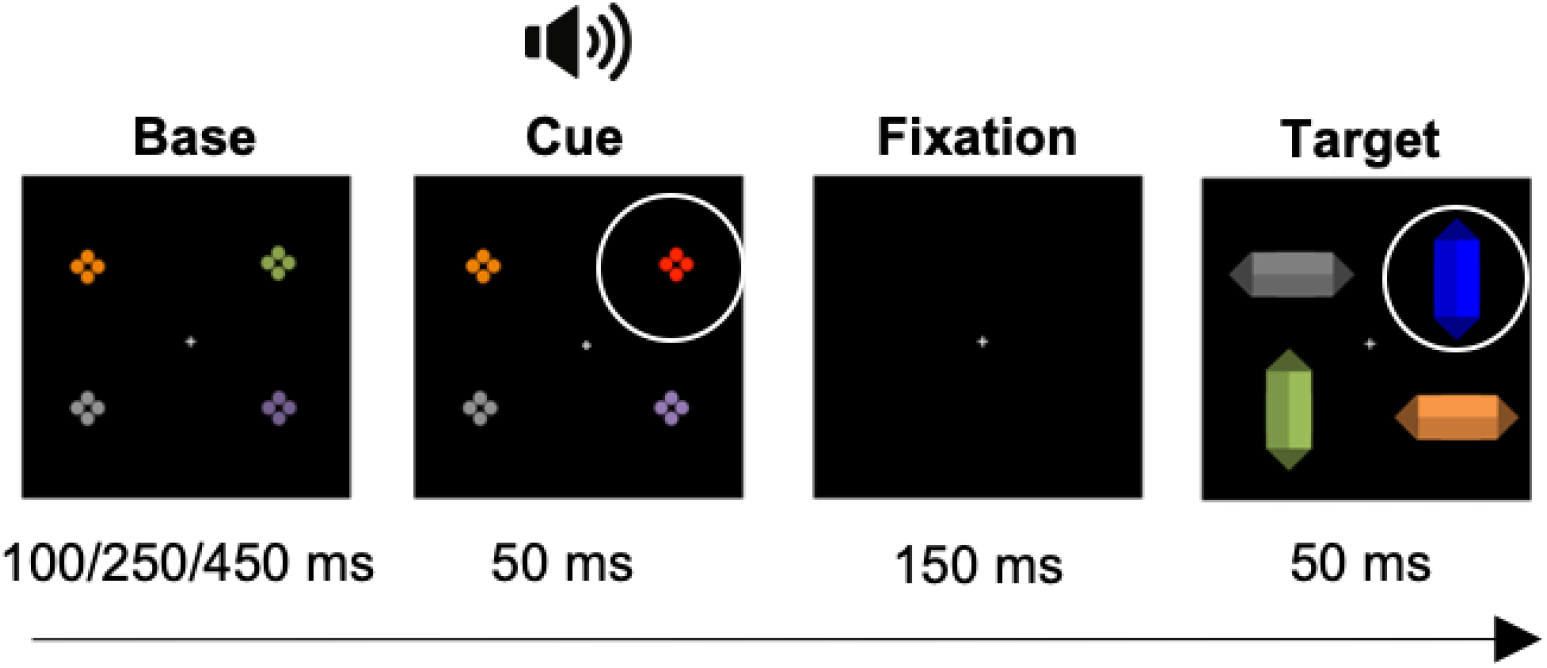
Experimental trial sequence for our paradigm. In this example, the blue target diamond is preceded by a nontarget-colour (NCC), i.e., red, ‘cue’, both highlighted here by white circles (that did not appear in the experimental task). A spatially diffuse sound was presented together with the onset of the colour change cue (on 50% of all trials), creating an audiovisual nontarget colour distractor (NCCAV).

Each experimental trial consisted of a sequence of arrays: base array, followed by cue array, followed by a fixation point, and finally a search array (**Error! Reference source not found.**). The base array contained four differently coloured sets of closely aligned dots, each dot subtending 0.1° × 0.1° of visual angle. Each set of dots could be one of four possible colours (according to the RGB scale): green (0/179/0), pink (168/51/166), gold (150/134/10), silver (136/136/132). Base array duration varied across trials (between 100, 250 and 450ms) to avoid building stimulus regularity-based predictions that could influence attentional control (Schwartze et al., 2011). In the cue array, one of the base array elements changed colour to either a target colour, or a nontarget colour that was not present in any of the elements before. The 2 cue colours were randomly selected with equal probability before each trial, and the colour change was not spatially predictive of the subsequent target location (same cue–target location on 25% of trials). On half of all trials, cue onset coincided with the onset of a pure sine-wave tone (2000Hz), presented from two loudspeakers on the left and right side of the monitor. Sound intensity was 80 dB SPL, as measured using a sound pressure meter as measured at the distance of the head using a CESVA SC-L sound pressure meter (CESVA, Barcelona, Spain). We set the time between the onset of the cue and the target to 200ms (cue array and fixation together) to reduce the likelihood of participants moving their eyes from the cue to the target, which would contaminate the data with eye movements (Yang et al. 2002).

Manipulating whether the distractor matched the target colour or not enabled us to measure task-set contingent attentional capture (TAC) across different age groups. This, in turn, provided insights into the age at which such visual attentional control mechanisms develop. In turn, manipulating the visual-only or audiovisual of the distractor allowed us to investigate the occurrence of multisensory enhancement of capture (MSE) in different age groups. This enabled us to investigate the development of attentional control processes engaged by task-irrelevant multisensory stimuli. The two above mentioned manipulations related to the cue resulted in two factors: Cue Colour (target colour-cue – TCC vs. nontarget-colour-cue – NCC) and Cue Modality (Visual – V vs. AudioVisual – AV), and, consequently, 4 experimental conditions: TCCV, NCCV, TCCAV, NCCAV.

The target array contained 4 elements (‘diamonds’) where 1 was always the colour-defined target. The targets and their preceding cues could have either a blue (RGB values: 31/118/220) or red (RGB values: 224/71/52) colour, and the target colour was counterbalanced across participants. The targets were given their diamond-like appearance by adding triangle shapes on the short sides of the bars and increasing and decreasing the luminance of certain sides of the bars by 20%.

Experimental sessions were conducted in a dimly lit, sound-attenuated room, with participants seated at a distance of 90 cm from a 23” LCD monitor with a resolution of 1080 × 1024 (60-Hz refresh rate, HP EliteDisplay E232). All elements were spread equidistally along the circumference of an imaginary circle against a black background, at an angular distance of 2.1° from a central fixation point. All visual elements were approximately equiluminant (~20cd/m^2^), as determined by a luxmeter placed at a position adjacent to participants’ eyes, measuring the luminance of the screen filled with each respective element’s colour. The means of three measurement values were averaged across colours and transformed from lux to cd/m^2^ in order to facilitate comparison with the results of Matusz & Eimer (2011).

To familiarise children with the task, a training block of 32 trials at 50% of regular task speed was administered. The subsequent full experimental session consisted of 8 blocks of 64 trials each, resulting in 512 trials in total. If participants did not respond within 5000ms of the target presentation, the next trial was initiated; otherwise the next trial was initiated immediately after a button press. Feedback on accuracy was given after each block, followed by a ‘progress (treasure) map’ which informed participants of the number of blocks remaining until the end, and during which participants could take a break and parents/caregivers could enter the testing room. To maintain motivation in younger participants, stickers on diamond-shaped sheets were offered during breaks following each session.

### 3. EEG acquisition and preprocessing

A 129-channel HydroCel Geodesic Sensor Net connected to a NetStation amplifier (Net Amps 400; Electrical Geodesics Inc., Eugene, OR, USA) was used to record continuous EEG data sampled at 1000Hz. Electrode impedances were kept below 50kΩ, and electrodes were referenced online to Cz. Offline filtering involved a 0.1 Hz high-pass and 40 Hz low-pass as well as 50 Hz notch and a second-order Butterworth filter (–12 dB/octave roll-off, computed linearly with forward and backward passes to eliminate phase-shift). Next, the EEG was segmented into peri-stimulus epochs from 100ms before cue onset to 500ms after cue onset. Epochs were then screened for transient noise, eye movements, and muscle artefacts using a semi-automated artefact rejection procedure. It has been noted previously that due to physiological differences between children and adults’ skulls and brains, these two groups require different artefact rejection criteria to prevent discarding clean EEG signal (Scerif et al., 2006; Shimi et al., 2015). Therefore, as in previous ERP research on developing populations (e.g., Melinder et al., 2010; Shimi et al., 2014b), we used an automatic artefact rejection criterion of ±100μV for adults and ±150μV for children, along with visual inspection. For children, additionally, only EEG data from trials with correct responses, and from blocks with over 50% accuracy were used, to fit behavioural data. Data from artefact contaminated electrodes across all groups were interpolated using three-dimensional splines (Perrin et al., 1987). Average numbers of epochs removed, and electrodes interpolated per participant in each age group can be found in Supplementary materials.

Cleaned epochs were averaged, baseline corrected to the 100ms pre-cue time interval, and re-referenced offline to the average reference. Next, due to persistent environmental noise present in the majority of the child and adult datasets even after initial filtering, an additional 50Hz notch filter was applied. All of the above steps were done separately for ERPs from the four cue conditions, separately for cues in the left and right hemifield. To analyse cue-elicited lateralised ERPs, single-trial data from all conditions with cues presented on the left were relabelled to have electrodes over the left hemiscalp represent activity over the right hemiscalp, and vice versa. After relabelling, the “mirror cue-on-the-right” single-trial data and the veridical “cue-on-the-right” data were averaged together, creating a single lateralised average ERP for each of the 4 cue conditions. As a result of this, we obtained 4 different ERPs, one for each of the 4 conditions (TCCV, NCCV, TCCAV, NCCAV). All preprocessing-and EEG analyses, unless otherwise stated, were conducted using CarTool software (available for free at www.fbmlab.com/cartool-software/; Brunet, Murray, & Michel, 2011).

### 4. Data analysis design

To reiterate, as we previously found both task-set contingent visual attention capture (TAC) and multisensory enhancement of attention capture (MSE) in adults (Matusz & Eimer, 2011), we used these as behavioural markers of top-down visual and bottom-up multisensory control processes. Next, we combined traditional N2pc component analyses with an electrical neuroimaging (EN) analytical framework.

#### 4.1. Behavioural analyses

Analyses were focused on reaction-time (RT) spatial cueing effects. This measure was derived by subtracting the mean RTs for trials where the cue and target were in the same location from the mean RTs for trials where the cue and target location differed, separately for each of the 4 cue conditions, following Matusz & Eimer (2011). Error rates were also analysed, in the form of percentages. Before the analysis, RT data were cleaned following a two-step procedure. First, blocks with mean accuracy below chance level (50%) were removed. Thus, in children, 15% of all blocks were removed (3% for 9-year-olds, 7% for 7-year-olds, and 37% for 5-year-olds respectively). In adults, all blocks were used due to high overall accuracy (>95%). Next, RT data from the remaining blocks was cleaned following the procedure of Gaspelin et al. (2015). Specifically, incorrect and missed trials were discarded, as were trials with RTs below 200ms and above 1000ms for adults, and below 200ms and above 5000ms for children. Moreover, all RTs above 2.5 *SDs* from individual participant’s mean RTs were also removed. Overall, 26% of all trials were removed (6% in adults, 28% in 9-year-olds, 29% in 7-year-olds, and 40% in 5-year-olds). Next, to verify if RT spatial cueing modulations were preserved after correcting for children’s general cognitive slowing, each individual’s RT’s per condition was divided by their average overall RT, and then converted to a percentage (following Gaspelin et al., 2015, see also Maylor & Lavie, 1998). ‘Raw’ and scaled RT data were normally distributed, and thus submitted to separate mixed-design four-way repeated-measures ANOVAs with one between-subject factor of Age (adults vs. 9-year-olds vs. 7-year-olds vs. 5-year-olds), and three within-subject factors: Cue Colour (target colour-cue -TCC vs. nontarget colour-cue -NCC), Cue Modality (Visual - V vs. AudioVisual - AV), and Cue-Target Location (Same vs. Different). Next, as part of follow-up analyses, data for each age-group were submitted to separate repeated-measures ANOVAs with within-subject factors: Cue Colour, Cue Modality, and Cue-Target Location. Error data were not normally distributed, and thus analysed using separate three-way Friedman tests for each child group, with factors Cue Colour, Cue Modality, and Cue-Target Location. In the case of adult control data, we conducted a three-way Durbin test instead, with factors Cue Colour, Cue Modality, and Cue-Target Location. All analyses, including post-hoc paired t-tests, were conducted using SPSS for Macintosh 26.0 (Armonk, NY: IBM Corp).

#### 4.2. Overview of ERP analyses

Given that the N2pc is a well-understood correlate of adult visual attentional control, we began our ERP analyses by conducting a canonical N2pc analysis, involving a comparison of the contralateral and ipsilateral average ERPs elicited across the 4 cue conditions. This way, we could compare the N2pc’s elicited by our visual and audiovisual distractors with the existing N2pc research in adults and children. Furthermore, those analyses helped us better bridge the previous and present canonical-analysis’ N2pc results with our electrical neuroimaging (EN; more details on the EN analyses below) analyses of the N2pc. Thus, we used data from the entire 129-channel electrode montage in our analyses of the contralateral versus ipsilateral ERP voltage gradients within an EN approach.

Since the aim of the current study was to identify the emergence of adult-like attentional control mechanisms in childhood, all ERP analyses in developmental groups followed a ‘normative’ framework. That is, the parameters typically used for canonical analyses of the N2pc in adult visual attention research were applied to children’s data. Below, we detail how we conducted the N2pc and EN analyses across age groups.

##### 4.2.1. Canonical N2pc analysis

We extracted the mean amplitude for each of the 4 cue conditions within a prescribed time-window, separately for the contralateral and the ipsilateral posterior electrodes. We used electrodes e65 and e90 – the EGI 129-channel equivalents of the canonical PO7/8 electrodes (e.g., Eimer et al., 2009; Kiss et al., 2008), and the time-window of 180-300ms post-stimulus onset (e.g., Luck & Hillyard, 1994; Eimer, 1996; Eimer 2014). We used these criteria to extract mean amplitudes for each of the 8 ERPs (4 cue conditions for ipsilateral and contralateral electrode each) for each of the four age groups. We then submitted these mean amplitude values to separate 3-way repeated measures ANOVAs, with within-subject factors: Cue Colour (TCC vs. NCC), Cue Modality (V vs. AV), and Contralaterality (Contralateral vs. Ipsilateral). For comparison, we also analysed the same data, choosing the electrode sites and time-window for extraction of mean amplitude values following a more data-driven approach (see Supplementary materials: Supplemental N2pc results).

##### 4.2.2. Electrical neuroimaging of the N2pc component

We have introduced EN analyses in the Introduction. Thus, here we provide more information on how EN has already been used to analyse mechanisms governing N2pc, how we designed our EN analyses of the N2pc in this study, as well as on the EN measures themselves.

Attentional control mechanisms can modulate the contralateral-to-ipsilateral gradients of voltage potentials across different conditions (see Matusz et al. 2019b). These gradients are not captured by canonical N2pc analyses, which only analyse the mean voltage difference between one contralateral and one ipsilateral electrode/electrode subset (out of a set of >20 or >100 electrodes). Further, the same mean voltage amplitude across two experimental conditions can arise from a completely different distribution of values across the scalp, and so, different sets of globally-distributed brain networks. However, traditional N2pc analyses cannot detect such differences (for tutorial-like demonstration, see Matusz et al., 2019b; Fig.3). The mechanism traditionally assumed to generate visual attentional effects in N2pc is a change in response strength within the same network (a “gain-control” mechanism), but canonical N2pc analyses cannot confirm or dispute this. EN analyses compensate for these limitations, as they take into account the entire scalp electrical field, and the measures it employs differentiate between network differences and response strength differences.

To analyse the global mechanisms underlying N2pc’s during visual control and multisensory control across development, we first computed a difference ERP where we subtracted the voltages over the ipsilateral hemiscalp from the voltages over the contralateral hemiscalp, separately for each of the 4 cue conditions. This resulted in a 59-channel lateralised ERP (without the uninformative midline electrodes from the 129-electrode montage). Next, this difference ERP was mirrored onto the other side of the scalp, creating a “fake” 129-electrode montage (with midline electrode values set to 0). It was on these mirrored “fake” 129-electrode lateralised difference ERPs that we performed the EN response strength and topography analyses, across the 4 cue conditions, for each of the 4 age groups.

###### 4.2.2.1. Strength-based modulations of the difference “N2pc-like” ERPs

We used Global Field Power (GFP) to investigate whether attentional modulations of cue-elicited lateralised ERPs resulted from differential response strength within statistically indistinguishable brain networks. GFP is a timepoint-to-timepoint standard deviation of voltage across the scalp, and can be plotted as a single waveform, just like any regular waveform. In an EN framework, differences in response strength in line with the gain-control account would be readily detected as GFP differences between experimental conditions over the N2pc time-window (Matusz et al., 2019b). To mirror our canonical N2pc analyses, we extracted the average GFP amplitudes in the 129-channel “fake” difference ERPs, measured across the canonical adult N2pc time-window of 180-300ms post-cue. We then submitted each age group’s 4 cue condition averages to separate 2 × 2 repeated-measures ANOVAs with Cue Colour (TCC vs. NCC) and Cue Modality (V vs. AV) as within-subject factors.

###### 4.2.2.2. Topographic modulations of the difference “N2pc-like” ERPs

Next, we investigated whether differences across the lateralised ERPs were driven by changes in electric scalp field topography, and in turn, changes in activated configurations of brain generators. To analyse topographic differences across conditions, we applied clustering to the group-averaged mirrored difference ERPs over their whole post-cue time course. Clustering can reveal periods of tens to hundreds of milliseconds of stable topographic activity, i.e., topographic “maps” (elsewhere referred to as “functional microstates”, e.g., Michel & Koenig, 2018). To this end, we used the hierarchical clustering method Topographical Atomize and Agglomerate Hierarchical Clustering (TAAHC), which, over a set of iterations, identifies configurations of clusters that explain certain amounts of global explained variance (GEV) in the ERP data (for more details see Matusz et al. 2019b; Murray et al., 2008). The optimal number of clusters is the smallest number of template maps accounting for the largest amount of GEV in the grand-averaged ERPs. To identify this number, we used the modified Krzanowski–Lai’s, the Cross Validation index, and the Dispersion criterion (Murray et al., 2008).

As part of our “normative” EN analyses, first we applied the TAAHC to the group-averaged *adult* ERP data and identified the optimal number of clusters that explain most of the adult ERP variance. Next, we tested how much the template maps seen in adults were present in the child groups’ ERPs, and how this involvement differed by age group. That is, for each age group separately, we investigated whether, and, if so, how strongly, each of the clusters identified in the adult group-averaged difference ERPs were present in the single-subject ERP developmental data (the so-called “fitting” procedure). Specifically, every time-point over the adult N2pc time-window in the cue-induced mirror difference ERPs of each tested child was labelled by the adult topographical map with which it best correlated spatially. The final output for each participant was the number of timeframes (in milliseconds) that each adult topographical map characterised the child’s ERP in the adult canonical N2pc time-window. These map durations were submitted to separate three-way 2 × 2 × 4 repeated-measures ANOVAs in each age group, with factors: Cue Colour (TCC vs. NCC) and Cue Modality (V vs. AV), and Map (Map1 vs. Map2 vs. Map3 vs. Map4) followed by post-hoc t-tests. Maps with durations under 10 contiguous timeframes were not included in the analyses. Greenhouse-Geisser corrections were applied where necessary to correct for violations of sphericity. Unless otherwise stated, map durations were reliably present across the time-windows of interest (statistically different from 0, as confirmed by post-hoc t-tests). Throughout the results, Holm-Bonferroni corrections were used to correct for multiple comparisons between map durations. Comparisons passed the correction unless otherwise stated.

## Results

### 1. Behavioural analyses

#### 1.1. ‘Raw’ reaction times

Mean RTs sped up progressively from 5-year-olds (1309ms) through 7-year-olds (1107ms) and 9-year-olds (836ms) to adults (594ms), which was reflected in a significant main effect of Age, *F*_(3, 127)_ = 94.7, *p* < 0.001, η_p_^2^ = 0.7. Here, 5-year-olds were reliably slower than 7-year-olds (*t*_(33)_ = 4.4, *p* < 0.001), who were slower than 9-year-olds (*t*_(32)_ = 5.7, *p* < 0.001), who were in turn, slower than adults (*t*_(32.5)_ = 5.1, *p* < 0.001). However, Age did not interact with any other factors (all *F*’s < 2, *p’*s > 0.1). Nonetheless, to adequately investigate differences between adults and children, and the developmental trajectory of attentional control processes, we analysed the raw RT data from each age group separately.

Firstly, in adults, there was a significant main effect of Cue Colour, *F*_(1, 38)_ = 36.9, *p* < 0.001, η_p_^2^ = 0.5, driven by faster responses on trials with target colour-cues (TCC, 607ms) than on trials with nontarget colour-cues (NCC, 618ms). Adults also showed generally faster responses on trials with sounds (AV, 605ms) than with no sounds (V, 620ms), *F*_(1, 38)_ = 76.1, *p* < 0.001, η_p_^2^ = 0.7. Overall behavioural capture effects in adults were reliable, i.e. responses were faster for trials where the cue and target location were the same (600ms) versus when they were different (624ms), *F*_(1, 38)_ = 110.9, *p* < 0.001, η_p_^2^ = 0.8. Further, as in the original Matusz and Eimer’s (2011) study, the adults’ overall behavioural capture effects differed depending on the colour of the cue, as shown by a 2-way Cue-Target Location × Cue Colour interaction, *F*_(1, 38)_ = 161.5, *p* < 0.001, η_p_^2^ = 0.8 (this is the TAC effect). This effect was driven by statistically significant behavioural capture effects for the TCC condition (48ms, *t*_(38)_ = 16.7, *p* < 0.001), but not the NCC condition (1ms, *t*_(38)_ = 0.2, *p* = 0.8; Figure 2, top left panel, and Figure 3 top left panel). Again, as in the original 2011 study, behavioural capture effects also differed when elicited by visual and audiovisual distractors, as shown by a two-way interaction between Cue-Target Location and Cue Modality, *F*_(1, 38)_ = 4.9, *p* =0.03, η_p_^2^ = 0.1 (this is the MSE effect). This effect was driven by larger behavioural capture effects elicited by AV (26ms, *t*_(38)_ = 10.8, *p* < 0.001) than by V cues (21ms, *t*_(38)_ = 8.9, *p* < 0.001; Figure 2, top left panel, and Figure 3 top left panel). The Cue Colour by Cue Modality interaction (*F* < 1) was not significant, and neither was the Cue-Target Location × Cue Colour × Cue Modality interaction (*F* < 3, *p* > 0.1). These results demonstrated that adults showed both reliable TAC and MSE in behaviour, replicating Matusz and Eimer (2011).

**Figure 2.**
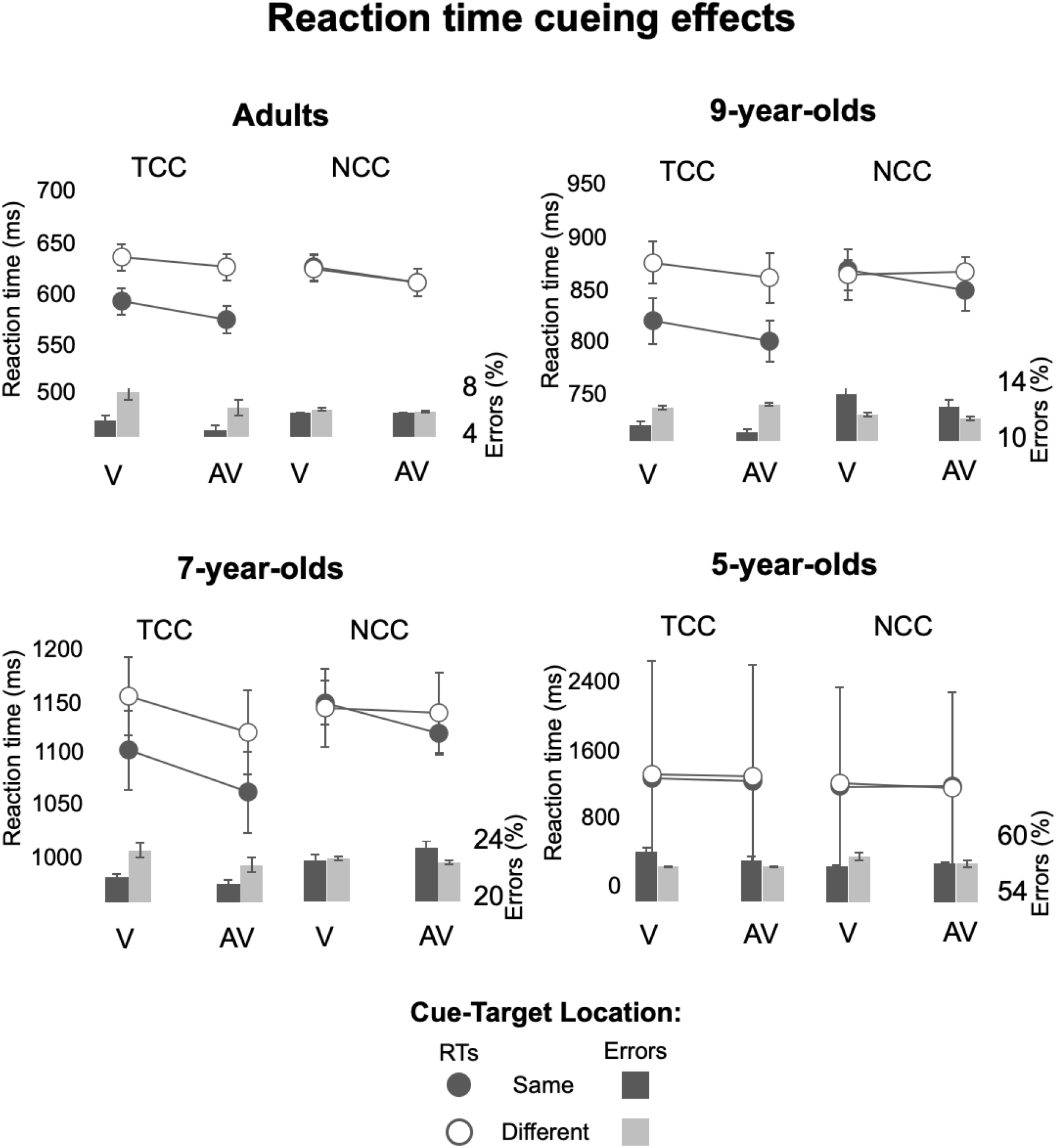
Mean reaction times shown for each of the 4 age groups on trials where Cue-Target Location was the same versus different, shown separately for target colour-cue (TCC) and nontarget colour-cue (NCC) trials, as well as visual (V) and audiovisual (AV) trials. Line graphs show the mean RTs, bar graphs show error rates (in percentages), and error bars represent the standard error of the mean. The RT ranges that best display the Spatial Cueing effects variability in the data are displayed. Thus, each age group’s scale has a different range, but the range lengths are the same (200ms), save for 5-year-olds where the variability was too large to maintain this range length.

**Figure 3.**
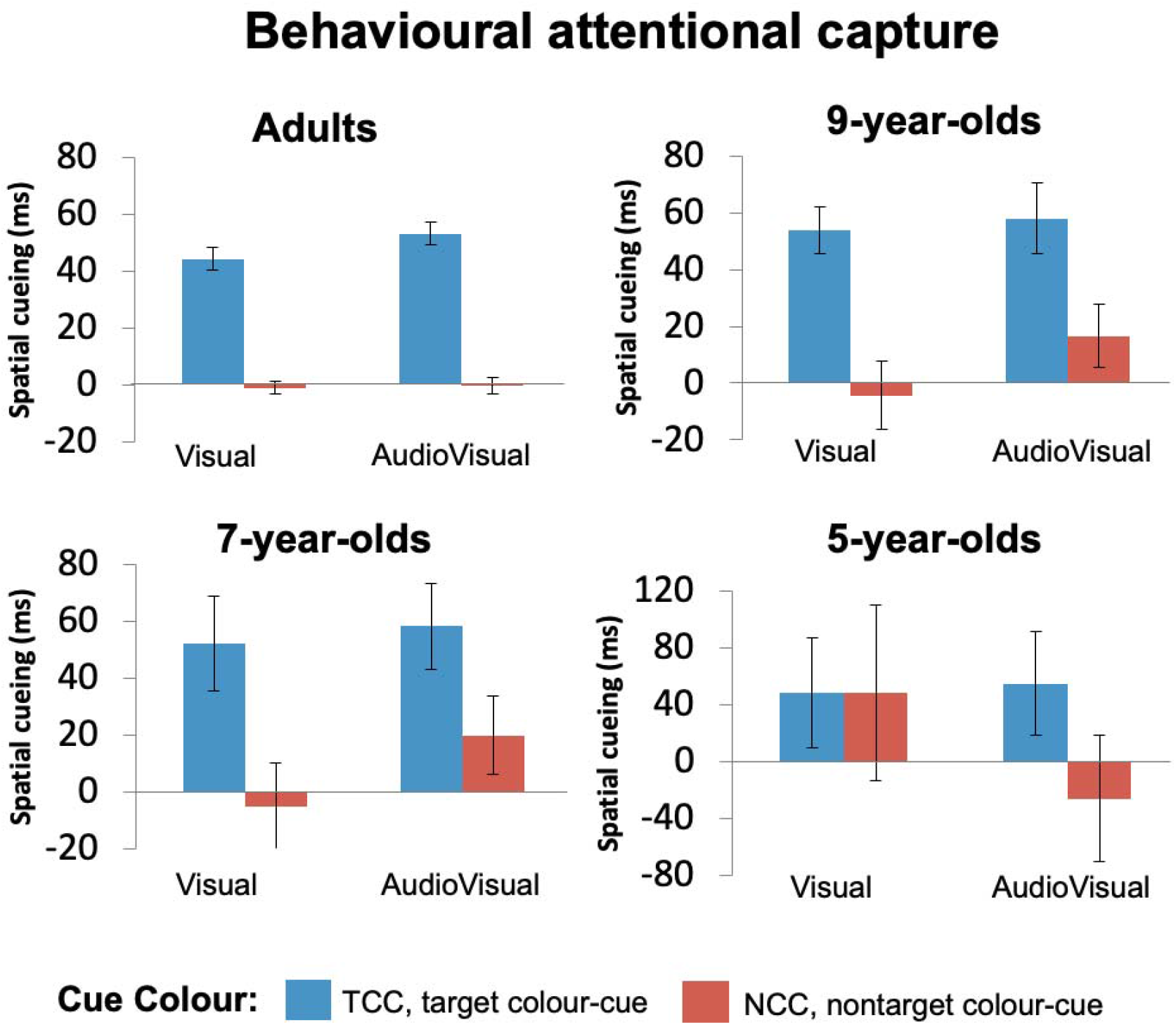
Bars coloured according to the figure legend in the image represent behavioural attentional capture indexed by mean RT spatial cueing effects, and error bars represent the standard error of the mean. Adults, 9-year-olds, and 7-year-olds all showed presence of top-down visual attentional control, exemplified by TAC. Specifically, all 3 age groups showed reliable attentional capture effects for target colour-cues, but not for nontarget colour-cues. In contrast, only in adults, attentional capture showed MSE.

Like adults, 9-year-olds responded faster on TCC trials (843ms) than on NCC trials (865ms), *F*_(1, 25)_ = 28.4, *p* < 0.001, η_p_^2^ = 0.5. Their overall behavioural capture effects were also reliable, with faster RTs for trials where the cue and target location were the same (839ms) versus when they were different (870ms), *F*_(1, 25)_ = 68.9, *p* < 0.001, η_p_^2^ = 0.7. Overall speeding up of responses on AV compared to V trials now showed the level of a nonsignificant trend (*F*_(1, 25)_ = 0.3, *p* = 0.08, η_p_^2^ = 0.1). However, the main question was whether behavioural capture effects in 9-year-old children would be modulated by the cues’ matching of the target colour, as well as the audiovisual nature of the cues. Notably, like in adults, 9-year-olds did indeed show TAC, as evidenced by a 2-way interaction between Cue-Target Location and Cue Colour, *F*_(1, 25)_ = 19.5, *p* < 0.001, η_p_^2^ = 0.4. This interaction was driven by significant capture effects for the TCC distractors (56ms, *t*_(25)_ = 8.3, *p* < 0.001), but not for the NCC distractors (6ms, *t*_(25)_ = 0.9, *p* = 0.7; Figure 2, top right panel, and Figure 3 top right panel). However, in contrast with adults, 9-year-olds did not show MSE, with no evidence for visually-elicited capture effects enlarged on AV vs. V trials, i.e., no 2-way Cue-Target Location x Cue Modality interaction, *F*_(1, 25)_ = 1.4, *p* = 0.3. Other interactions failed to reach statistical significance (All *F*’s < 2, *p’*s > 0.1). With this, we can conclude that 9-year-olds showed reliable TAC, but not MSE, in behaviour.

In 7-year-olds, like in adults, responses were faster for trials with TCC cues (1112ms) than for NCC cues (1138ms), *F*_(1, 37)_ = 18.7, *p* < 0.001, η_p_^2^ = 0.3, and were also faster for trials with AV cues (1111ms) than V cues (620ms), *F*_(1, 37)_ = 8.6, *p* = 0.006, η_p_^2^ = 0.2. Further, overall capture effects were again reliable, with faster responses on cue-target location same (1109ms) versus different (1140ms) trials, *F*_(1, 37)_ = 14, *p* < 0.001, η_p_^2^ = 0.4. Just as in the two older groups, 7-year-olds, did show TAC, as shown by a Cue-Target Location x Cue Colour interaction, F = 6.4, *p* = 0.02, η_p_^2^ = 0.2. This was driven by reliable cueing effects elicited by TCC distractors (55ms, *t*_(37)_ = 4.8, *p* < 0.001), but not by NCC distractors (7ms, *t*_(37)_ = 0.6, *p* = 1; Figure 2, bottom left panel, and Figure 3 bottom left panel). However, as in 9-year-olds, 7-year-olds’ visually-induced attentional capture effects did not show MSE, with no 2-way Cue-Target Location x Cue Modality interaction failing to reach significance, *F*_(1, 37)_ = 2.1, *p* = 0.2. Other interactions also did not reach statistical significance (All *F*’s < 2, *p*’s > 0.1). It thus appeared that 7-year-olds, like 9-year-olds before them, showed reliable TAC, but not MSE.

In 5-year-olds, as in the other age groups, we observed reliable overall attentional capture effects *F*_(1, 27)_ = 14, *p* < 0.001, η_p_^2^ = 0.4, driven by faster responses for cue-target location same (1312ms) versus different (1343ms) trials. However, there was no evidence for either of the two key interactions, specifically, the Cue-Target Location x Cue Colour interaction (*F*_(1, 27)_ = 1.4, *p* = 0.2), or the Cue-Target Location x Cue Modality interaction (*F*_(1, 27)_ = 0.4, *p* = 0.5). In further contrast with the older age groups, overall RTs were not affected by the colour of the cue, as shown by a nonsignificant main effect of Cue Colour, *F*_(1, 27)_ = 2.6, *p* = 0.1. In one final contrast, faster responses on AV versus V trials showed only a nonsignificant trend, *F*_(1, 27)_ = 3.5, *p* = 0.07, η_p_^2^ = 0.1. No other interactions reached statistical significance (All *F*’s < 2, *p*’s > 0.1). The 5-year olds, therefore, did not show reliable TAC nor MSE in behaviour.

#### 1.2. RTs corrected for children’s cognitive slowing

Even after correcting for children’s overall cognitive slowing, all child groups showed the same patterns of results as in the raw RT analyses. That is, 9-year-olds and 7-year-olds showed TAC but not MSE, and 5-year-olds did not show either effect. The full description of the results on data corrected for slowing can be found in Supplementary materials.

#### 1.3. Error rates

Since error data were not normally distributed, we conducted a 1-way Kruskal–Wallis *H* test to test for differences between groups, and 3-way Friedman tests (or Durbin tests where there were no errors for a given condition) to test for differences within each age-group. Overall, error rates were highest in 5-year-olds (57%), and steadily reduced in 7-year-olds (23%), followed by 9-year-olds (12%), and adults (6%), χ^2^(3) = 81.4, *p* < 0.001. In adults, error rates were modulated by Cue-Target Location χ^2^(1) = 8.7, *p* = 0.003, such that fewer errors were made on trials where the cue and target location was the same (5.5%) than when they were different (6.6%). Error rates were not significantly modulated by Cue Colour or Cue Modality (all *p*’s > 0.1). In 9-year-olds, 7-year-olds, and 5-year-olds alike, error rates were not significantly modulated by Cue-Target Location, Cue Colour or Cue Modality (all *p*’s > 0.1).

### 2. ERP analyses

#### 2.1. Canonical N2pc analysis

In adults, there was a reliable overall N2pc, as demonstrated by a statistically significant main effect of Contralaterality, *F*_(1, 38)_ = 17.8, *p* < 0.001, η_p_^2^ = 0.3, where the mean contralateral amplitude (−0.4μV), was larger than the ipsilateral amplitude (0.1μV). Consequently, the contra-ipsilateral difference had a mean overall amplitude of −0.5μV. As expected, cue-elicited N2pcs’ differed in their magnitude depending on the cue colour, as supported by a Contralaterality x Cue Colour 2-way interaction, *F*_(1, 38)_ = 17, *p* < 0.001, η_p_^2^ = 0.3. This interaction was driven by a reliable N2pc for target colour-cues (−0.69μV; **Error! Reference source not found**. Figure 4, first panel, top and bottom left windows) but not for nontarget colour-cues (−0.25μV; Figure 4, first panel, top and bottom right windows). This result demonstrated presence of TAC in adult N2pc’s. However, there was no evidence for MSE, as there was no Contralaterality x Cue Modality 2-way interaction (*F* < 1).

**Figure 4.**
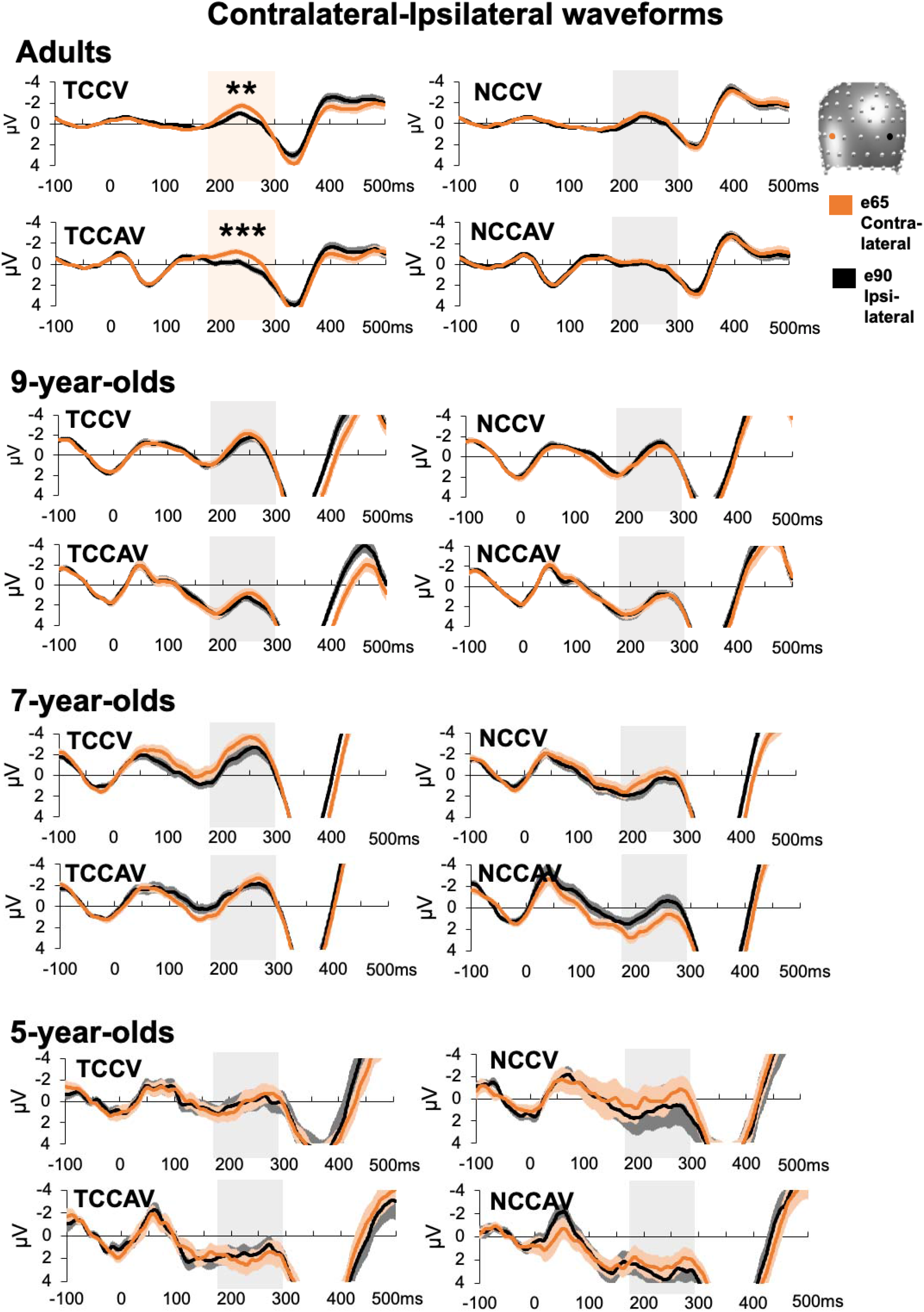
N2pc waveform results. Mean amplitude values are shown at contralateral and ipsilateral electrode sites, indicated in orange and black, per the head model and legend on the figure. The N2pc time-window of 180-300ms is highlighted in light orange, where the contra-ipsi difference is significant, and light grey where it is not. Significance levels are denoted as follows: ** < .01, *** < .001. Adults show significant contra-ipsi differences, that is reliable N2pc’s, for target-colour cues (TCC) but not nontarget colour-cues (NCC). In children, there was no reliable N2pc in any of the four conditions.

Interestingly, the N2pc amplitudes elicited by TCC and NCC cues were modulated by sound presence, as shown by a 3-way interaction between Contralaterality, Cue Colour, and Cue Modality, *F*_(1, 38)_ = 8, *p* = 0.007, η_p_^2^ = 0.2. We first analysed this interaction as a function of Cue Modality. First, for AV cues, mean N2pc amplitudes elicited by TCCAV were larger (−0.8μV) than mean amplitudes for NCCAV cues (−0.2μV), *t*_(38)_ = 5, *p* < 0.001. In contrast, for V cues, there was no statistically significant difference in mean N2pc amplitudes elicited by NCCV cues (−0.3μV) and TCCV cues (−0.6μV), *t*_(38)_ = 1.8, *p* = 0.2. When we analysed the 3-way interaction as a function of Cue Colour, for both TCC and NCC distractors, differences in mean N2pc amplitude between AV and V were at the level of a nonsignificant trend (*t*_(38)_ = 1.8, *p* = 0.06, and *t*_(38)_ = 1.4, *p* = 0.07, respectively). Other effects did not reach statistical significance (All *F*’s < 1), except the main effects of Cue Colour, *F*_(1, 38)_ = 8.4, *p* = 0.006, η_p_^2^ = 0.2 (driven by larger ERP amplitudes for TCC −0.3μV, than for NCC −0.03μV, and Cue Modality, *F*_(1, 38)_ = 7.1, *p* = 0.011, η_p_^2^ = 0.2 (driven by larger ERP amplitudes for V, −0.3μV, than for AV, 0.06μV). Thus, although MSE was not observed in N2pc’s, adult’s overall ERP data was jointly modulated by visual and multisensory attentional control. This effect seemed to be driven by reliable difference between TCC and NCC distractors on trials where distractors were AV but not V.

For the child age groups, 2 × 2 × 2 repeated-measures ANOVAs were conducted on mean amplitude values from adult electrodes over the adult time-window. In no child group was there a significant main effect of Contralaterality (9-year-olds: *F*_(1, 25)_ = 0.4, *p* = 0.6; 7-year-olds: *F*_(1, 37)_ = 0.04, *p* = 0.8; 5-year-olds: *F*_(1, 27)_ = 0.2, *p* = 0.6; Figure 4, 2^nd^, 3^rd^, and 4^th^ panels), and therefore, no N2pc. For this reason, we will not report other results unrelated to Contralaterality (they are available in Supplementary Materials: Supplemental N2pc results). To rule out the possibility that a lack of effects in children was due to literature-based values being suboptimal, we conducted an additional analysis where the N2pc time-window and electrode sites were selected from the adult data in a more data-driven fashion. We report the details of the procedure and results in Supplementary Materials. Crucially however, this approach also showed no significant main effect of Contralaterality (All *F*’s < 1), and thus no presence of an N2pc.

#### 2.2. Electrical neuroimaging of the N2pc component

An ANOVA on the average GFP values per condition revealed no significant main effects or interactions in adults, 9-year-olds, 7-year-olds, or 5-year-olds (All *F*’s < 1). Full results can be found in Supplementary Materials: Supplemental GFP results. For graphical representations of the GFP results, we direct the reader to Supplemental Figure 1 in the Supplementary Materials.

The segmentation of the post-cue period of the adult data revealed 9 clusters which explained 82.8% of the GEV in the group-averaged ERPs. We remind the reader that topographical analyses were conducted on difference ERPs, which accounts for the lower rates of GEV. Next, a fitting procedure on the adult single-subject data revealed 4 template maps which characterised the N2pc time-period of 180-300ms post-cue. A 2 × 2 × 4 ANOVA on the mean durations of the 4 maps identified in the adult data revealed a main effect of Map, *F*_(3, 114)_ = 18.3, *p* < 0.001, η_p_^2^ = 0.3, where Map4 predominated (i.e. had the longest duration of all maps) the N2pc time-window across conditions (Figure 5, middle left panel). This demonstrated that adults had stable patterns of lateralised ERP activity. Hereafter, we did not follow up the main effect of Map with post-hoc tests, as it was not informative as to the presence of TAC or MSE in topography.

**Figure 5.**
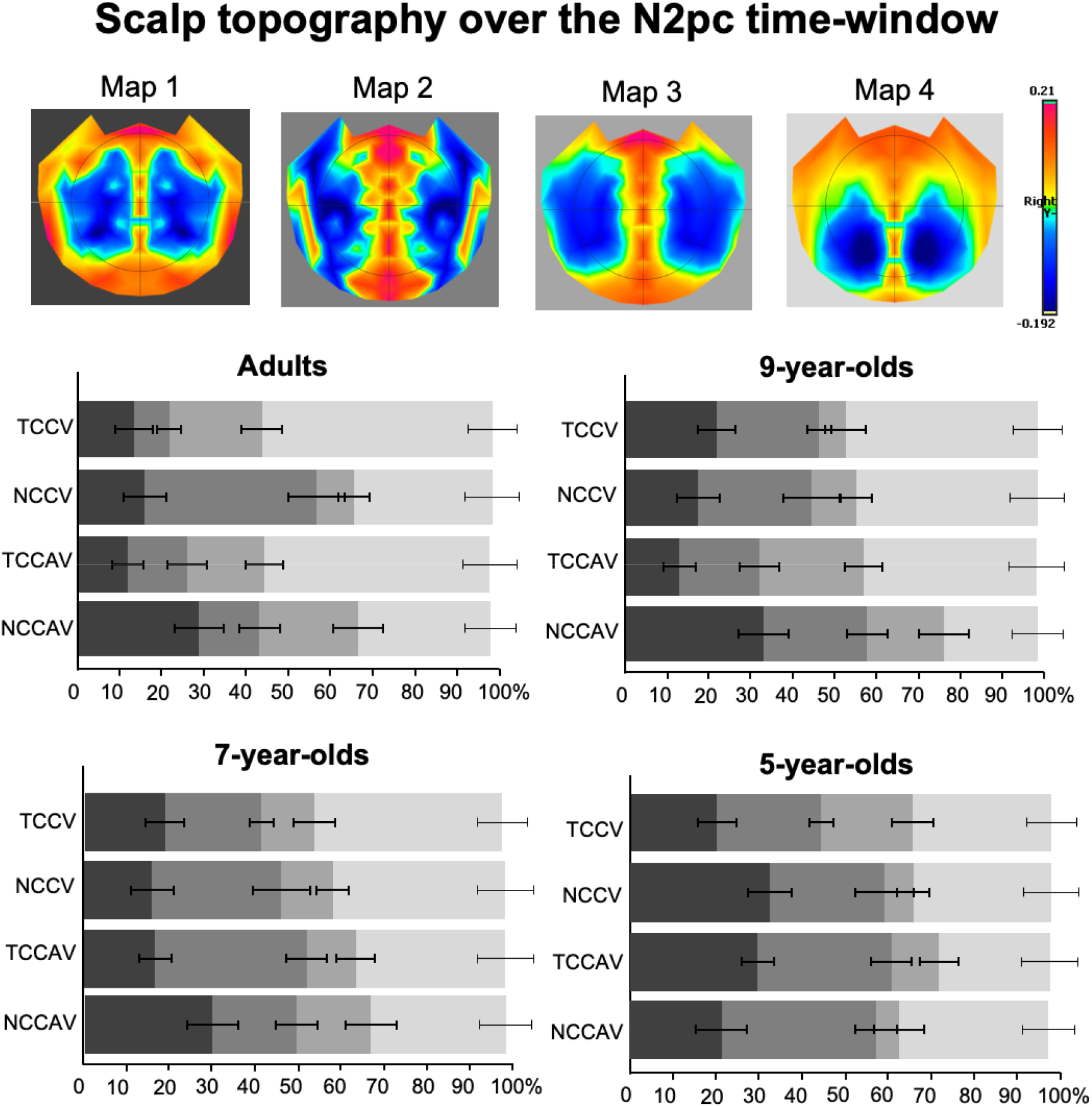
Scalp topography of the 4 lateralised difference template maps elicited over the N2pc time-window as a function of cue condition and observer age group. The four template maps resulting from the segmentation of the adult lateralised ‘mirrored’ difference ERP data are shown in the upper row. The bar graphs below represent each difference template map’s relative duration (% ms) over the N2pc time window, shown separately for the adults and the 3 younger groups, and for each of the V and AV cue conditions separately. Bars in the graphs are coloured according to their map’s backgrounds in the top row, and error bars denote the standard error of the mean. As visible in the lower graphs, Map 4 was the most dominant in adults, 9-year-olds, and 7-year-olds, while 5-year-olds did not have a clear map dominance pattern. Only in adults’ duration of Map 4 was modulated by cue type that is whether cue colour matched that of the target-colour.

There was a 2-way interaction between Map and Cue Colour, *F*_(2.4, 89.1)_ = 12, *p* < 0.001, η_p_^2^ = 0.2. Following up this interaction by the factor of Cue Colour showed that Map4 predominated responses to TCC (67ms) compared to NCC distractors (40ms), *t*_(38)_ = 5.2, *p* < 0.001, while Map2 predominated responses to NCC (34ms) compared to TCC distractors (13ms), *t*_(38)_ = 3.9, *p* = 0.004. Other maps did not differ significantly between TCC and NCC cues (all *p*’s > 0.1). Hereafter, differences in map predominance that are not reported here were not statistically significant (*p*’s > 0.1). Following up the interaction by the factor of Map revealed that for TCC cues, Map4 (67ms) overall predominated the N2pc time-window compared to all other maps – Map1 (15ms), *t*_(38)_ = 7.7, Map2 (13ms), *t*_(38)_ = 8, and Map3 (25ms), *t*_(38)_ = 6.3, all p’s < 0.001, while no map differed in their predominance of responses to NCC distractors (all *p*’s > 0.1). These results suggest that Map4 drove the processing of TCC distractors, while no particular map was more implicated than others in the processing of NCC distractors. Finally, the map modulations by Cue Colour demonstrated here support the presence of TAC in adult ERP topography. Thus, it appeared that the Map x Cue Colour interaction was driven by modulations of Map2 and Map4 presence for different cue colours, where Map4 is especially implicated in the processing of target-colour cues.

In contrast to canonical N2pc analysis results, topographic map presence over the N2pc time-window interacted with Cue Modality, as evidenced by a 2-way interaction, *F*_(3, 114)_ = 3.2, *p* = 0.027, η_p_^2^ = 0.1. A follow-up by Cue Modality revealed that Map2 predominated responses when cues were purely visual (V, 30ms) than audiovisual (AV, 17ms), at the level of a nonsignificant trend, *t* = 2.8, *p* = 0.08. However, a follow-up by Map revealed that Map4 predominated the N2pc time-window compared to other maps for both AV cues (Map4 [53ms] vs. Map1 [25ms], *t*_(38)_ = 4.3, Map2 [17ms], *t*_(38)_ = 5.6, Map3 [26ms], *t*_(38)_ = 4.2, all *p*’s < 0.001), and for V cues (Map4 [54ms] vs. Map1 [18ms], *t*_(38)_ = 5.7, Map2 [30ms], *t*_(38)_ = 3.7, Map3 [19ms], *t*_(38)_ = 5.5, all *p*’s < 0.001). Taken together, it appeared that Map2 may be implicated more strongly in topographic modulations of lateralised ERPs by Cue Modality, whereas Map4 was the main map driving the processing of both AV and V cues.

Finally, the 3-way Map x Cue Colour x Cue Modality interaction was significant, *F*_(3, 114)_ = 5.4, *p* = 0.002, η_p_^2^ = 0.1. When followed up as a function of Cue Colour, for NCC distractors, Map2 predominated responses to V cues (50ms) over AV cues (18ms), *t*_(38)_ = 4.7, *p* < 0.001. Yet, for TCC distractors, all map durations were comparable between V and AV cues (all *p*’s > 0.1). Next, when following up as a function of Cue Modality, for AV cues, Map4 predominated responses to TCC distractors (67ms) compared to NCC distractors (40ms), *t*_(38)_ = 3.8, *p* = 0.004. Likewise, for V cues, Map4 predominated TCC (67ms) than NCC (39ms) distractor responses, *t*_(38)_ = 3.6, *p* = 0.003. However, Map2 also predominated responses to NCC (50ms) over TCC distractors (10ms), for V cues, *t*_(38)_ = 5.4, *p* < 0.001. Thus, maps that are sensitive to TAC and MSE appear to interact, suggesting that top-down visual attentional control and bottom-up multisensory attentional control may share neural generators.

To explore if and when the above adult topographical EEG patterns are present in children, we submitted each child age-groups’ data within the 180–300ms time-window to a fitting procedure, where child topographical data were labelled according to the adult template maps with they which they best correlated spatially.

For 9-year-olds, the ANOVA revealed a main effect of Map, *F*_(3, 75)_ = 9.2, *p* < 0.001, η_p_^2^ = 0.3, and, like in adults, Map4 predominately characterised ERPs during the N2pc time-window (Figure 5, middle right panel). Map presence was modulated only by Cue Modality, as evidenced by a 2-way interaction between Map and Cue Modality, *F*_(3, 75)_ = 3.4, *p* = 0.04, η_p_^2^ = 0.1. A follow up by Cue Modality found that Map3 predominated the responses to AV (27ms) compared to V cues (11ms), *t*_(25)_ = 2.6, *p* = 0.02, while Map4 predominated responses to V (55ms) compared to AV cues (39ms), *t*_(25)_ = 2.5, *p* = 0.02. However, the map that was sensitive to the (audio)visual nature of the cues in adults, Map2, was comparable in how it predominated responses to V cues (31ms) and AV cues (27ms), *t*_(25)_ = 0.7, *p* = 1. In a follow-up as a function of Map, there were no significant differences between map predominance for AV cues (all *p*’s > 0.1). For V cues, however, Map4 (55ms) predominated overall, compared to all other maps (Map1 [24ms], *t*_(25)_ = 4, Map2 [32ms], *t*_(25)_ = 3.7, Map3 [11ms], *t*_(25)_ = 5.8, all *p*’s < 0.001). In a marked contrast to adults, 9-year-olds did not show the other 2-way interaction of interest, Map x Cue Colour (*F*_(3, 75)_ = 1.3, *p* = 0.3). Other interactions also failed to reach statistical significance (all *F*’s < 2, *p*’s > 0.1). Taken together, 9-year-olds seemed to show adult-like MSE (a Map x Cue Modality 2-way interaction). Even though they did not show a modulation of the adult MSE-sensitive map, 9-year-olds’ overall topographic results were like those of adults, with a predominance of Map4 across conditions.

In 7-year-olds, there was also a main effect of Map, *F*_(2.3, 85.5)_ = 9.7, *p* < 0.001, η_p_^2^ = 0.2, with a predominance of Map4, akin to the two older age groups (Figure 5, bottom left panel). Unlike in older age groups, however, no other main effects or interactions reached statistical significance (all *F*’s < 3, *p*’s > 0.1). This included the 2-way interactions of interest, Map x Cue Colour (*F*_(3, 111)_ = 0.7, *p* = 0.6) and Map x Cue Modality (*F*_(2.4, 87.3)_ = 1.3, *p* = 0.3). We can therefore conclude that 7-year-olds’ topography did not show adult-like TAC or MSE, although their overall topographic pattern could be considered adult-like.

Finally, 5-year-olds also showed a main effect of Map, *F*_(3, 81)_ = 6.3, *p* < 0.001, η_p_^2^ = 0.2, but here, there was no clear map predominance pattern (Figure 5, bottom right panel). No other main effects or interactions reached statistical significance (All *F*’s < 1), including the two 2-way interactions of interest, Map x Cue Colour (*F*_(2.1, 57)_ = 0.8, *p* = 0.4) and Map x Cue Modality (*F*_(2.3, 61.6)_ = 0.7, *p* = 0.5). With this, 5-year-olds seemed not to show adult-like TAC, MSE, or overall pattern of map presence.

## Discussion

In cluttered learning environments like classrooms, children must focus their attention on relevant information and ignore unimportant information. There is modest research on the differences in neuro-cognitive mechanisms governing visual (and less so, auditory) attentional control between adults and children. In contrast, little is known about the development of multisensory attentional control mechanisms. Our study aimed to clarify how adult-like visual and multisensory attentional control mechanisms develop side by side over the course of primary education. We tested this by combining traditional behavioural and ERP measures of attentional selection with multivariate electrical neuroimaging analyses.

### 1. Developmental trajectory of visual attentional control

Behaviourally, we replicated in adults both task-set contingent visual attention capture (TAC) and multisensory enhancement of attention capture (MSE). We did so in a larger sample and with small adjustments to the Matusz and Eimer (2011)’s paradigm (to make the paradigm more child friendly). Children as young as 6-7 years (as well as 8-9-year-olds) showed adult-like magnitudes of both facilitatory and inhibitory visual attentional control. Specifically, they showed large spatial cueing effects in response to target-colour cues, and null cueing effects to nontarget-colour cues, respectively. These effects held even after correcting for children’s overall slower processing speed. This finding suggests that children may reach an adult-like state of visual feature-specific attentional control like TAC already at the age of 6-7 years. Other studies have corroborated this. For example, Oh-Uchi et al. (2010) found that the magnitude of attentional capture elicited by nontarget colour singleton stimuli was comparable between adults and 6-year-olds (albeit the study did not account for developmental differences in the overall RTs). Additionally, Greenaway and Plaisted (2005) showed, in a replication of the original Folk et al. (1992) study, colour distraction already in 11-year-olds.

Behavioural findings were extended by EN findings. In adults, topographical ERP analyses revealed two stable patterns of brain activity (template maps) that were each modulated by TAC and by MSE. Interestingly, the adult TAC-sensitive template map dominated the N2pc time-window overall -in adults, 7-, and 9-year-olds. However, in the child groups, the predominance of the adult TAC-sensitive map was not modulated by target-colour-matching, as those groups did not show evidence for a Map x Cue Colour interaction. Nonetheless, the child groups showed adult-like visual attentional control in behaviour also as well as the recruitment of brain networks modulated by distractor colour in adults. This at least indirectly supports that already 7-year-olds can deploy their top-down attention in a way that could be considered adult-like.

Our youngest group, 5-year-olds, did not show a reliable TAC effect. This result contrasts with the only other study on TAC in young children (Gaspelin et al. 2015), which found TAC in young children, albeit to a smaller degree than in adults. This finding is even more surprising considering that children in their study were younger (4.2 years) than in ours (5 years). The difference in the results may arise from the fact that Gaspelin et al. (2015) used colour singleton cues, rather than non-salient colour changes cues, which likely facilitated their propensity to capture attention. Additionally, young participants in our study could have been affected by heightened discomfort and fatigue due to the addition of fully irrelevant sound stimuli in our paradigm or the concurrent EEG recording that increased the total testing time. We nevertheless provided new findings regarding visual attentional control in such young children. Namely, 5-year-olds effectively utilised the non-salient colour-change distractors to orient their spatial attention, and these effects were found despite the large variability in this group’s RTs. This idea is further supported by our EN analyses that revealed in 5-year-olds the stable spatially selective (and so indicative of attentional selection in space) patterns of EEG activity observed in adults. This result is novel and important as it suggests that developed (adult) and nascent (young children’s) top-down visual control are instantiated at least partly through similar neuro-cognitive mechanisms. Additionally, as 5-year-olds in our study were relatively familiar with the school context, it is tempting to interpret these results as being driven by schooling experience acting as training of young children’s attentional control. At 5 years, Swiss children learn how to interact appropriately with peers and teachers and receive training in foundational skills such as phonics and numerical awareness (CIIP, 2012). Thus, by age 6-7, they have been in formal education for two years, and studies such as, for example, Brod et al. (2017), have shown evidence that even one year of formal schooling experience can improve attentional control. However, to provide evidence for a direct role of schooling in the early emergence of adult-like visual top-down control, one would likely need a similar approach to Brod et al.’s involving comparing 5-year-olds who entered first grade and those who remained in the kindergarten.

### 2. Development of attentional control processes engaged by multisensory stimuli

Despite adapting the original paradigm to children and adding an EEG measurement, also adults in our study showed the behavioural MSE effect. This corroborates the particular salience of multisensory distractors (Santangelo & Spence 2007; Talsma et al. 2010; van der Burg et al. 2011; Matusz & Eimer 2011; Matusz et al. 2015; 2019a; Turoman et al. 2020a), further supported by evidence that multisensory integration can occur at stages of brain processing preceding those influenced by top-down processes (Giard & Peronnet 1999; Cappe et al. 2010; reviewed in Talsma et al. 2010; De Meo et al. 2015; Murray et al. 2016; ten Oever et al. 2016). Surprisingly, none of the children groups showed MSE in behaviour, even though children supposedly have weaker attentional control than adults (Bunge et al. 2002; Hwang et al. 2010) and so should be theoretically more sensitive to more salient distractors. The null MSE effects in children could potentially be explained by the slow maturing of multisensory processes. Till now, this protracted development was shown mainly for attended, task-relevant objects (Gori et al., 2008; 2012; Barutchu et al. 2009; Denervaud et al. 2020). Our results would extend this principle to task-*irrelevant* objects. Notably, however, we did not study multisensory integration per se, but rather cross-modal audio-visual interactions, which should be present already at age 5 (e.g. Bahrick, 2001; Petrini et al. 2015; Broadbent et al., 2018). Therefore, the null developmental MSE effects in our study might have arisen from a combination of strong task demands (i.e. paying attention to fast-disappearing targets of a particular colour embedded in an array of similar coloured shapes) and cross-modal audiovisual interactions within task-irrelevant objects that may be less automated in children. Finally, the behavioural MSE itself is not large even in adults, ranging between 5 and 10ms (see also Matusz & Eimer 2011; Turoman et al. 2020a). Notwithstanding, using EN analyses we revealed the sensitivity of children’s brains to the multisensory nature of distractors and the recruitment of adult-like neuro-cognitive mechanisms for this purpose, from 8 years onwards. In other words, from the age of 8-9 years multisensory processes can permeate goal-directed behaviour even when gauged by task-irrelevant objects. Additionally, our EN analyses revealed that multisensory distraction activates spatially-selective brain mechanisms, which contrasts with previous, rare findings (van der Burg et al. 2011). Together, our results demonstrate the EN analyses are sensitive measures capable of revealing brain and cognitive mechanisms that may not be readily visible with behavioural measures or traditional ERP analyses.

Our multisensory developmental results could have important applied implications. First, they highlight the potential benefits of largely involuntary multisensory processes for attending to and encoding objects and symbols into memory, thus extending the known important role of top-down *visual* attention for learning and memory to multisensory attention processes (e.g., Astle & Scerif 2011; Shimi & Scerif, 2017). Second, our findings indicate that classroom design could benefit from minimising the risks of multisensory distraction (for detrimental effects of *unisensory* distraction on learning see: Fisher et al., 2014; Massonnié et al., 2019). Finally, our results demonstrating the earlier development of top-down unisensory attention than bottom-up multisensory attention could help better tailor brain rehabilitation and sensory-substitution training programs to participants’ age (e.g. Murray et al. 2015; Matusz et al. 2018; Buchs et al. 2019).

### 3. The N2pc as a marker of developing real-world attentional control

In adults, canonical N2pc analyses showed TAC, which mirrored the behavioural results and replicated visual attention research (e.g., Kiss et al. 2008; Eimer et al. 2009). However, N2pc results did not mirror behavioural MSE. This was not too surprising as the only other comparable study (van der Burg et al. 2011) showed weak evidence for attentional capture by multisensory distractors in N2pc. With this, while recording N2pc to distractors may be valuable for investigating visual attentional control in laboratory settings, our results suggest that the validity of the N2pc may be limited when studying attentional control in naturalistic, multisensory, settings.

In children, no adult-like N2pc’s were found in response to visual or audiovisual distractors in the canonical N2pc analyses. This contrasts with extant visual developmental studies, where a delayed but significant N2pc had been reported by the age of 9 (Couperus & Quirk, 2015; Shimi et al., 2015; see Sun et al. 2018 for N2pc in 9-15-year-olds). However, those studies recorded N2pc’s to targets, whereas, in order to test for TAC and MSE effects, we recorded them to distractors. Thus, one potential explanation for null N2pc’s in our children groups is that any distractor-elicited N2pc, which may arise more slowly than in adults, were overshadowed by the responses related to the target, which appeared already 200ms after the distractor onset. Had our analyses stopped at the N2pc, one could have concluded that attentional control processes like TAC and MSE are simply not elicited in children. However, with the use of EN, we revealed that adult-like spatially-selective brain mechanisms (that are captured partly by canonical N2pc analyses) were present at age 7 onwards, corroborating our behavioural results. To our knowledge, this is the youngest age group ever in which such spatially-selective -N2pc-like -brain mechanisms effects have been reported (cf. Couperus & Quirk, 2015; Shimi et al. 2015). Notably, with our approach, we revealed adult-like spatially-selective patterns for visual top-down and bottom-up multisensory attention that are at least partly independent. Here, the overall adult-like predominance of one map over others (Map4) was present already in 7-year-olds.

Our findings challenge the idea of the canonically analysed N2pc as a viable general marker of attentional selection; the N2pc’s failed to show sensitivity of adult attentional control to multisensory distractors and to any distractors in the child groups. However, when combining the N2pc and EN analyses, that is, taking into account whole-brain activity over the N2pc time-period, we obtained neurophysiological markers of 1) the previously elusive sensitivity of visual attentional selection to bottom-up multisensory processing, and 2) adult-like attentional control processes in children as young as 7. The second finding is certainly promising for the use of EN to study the development, of attentional control and beyond. Developmental studies compare adults and children in their attentional skills (e.g., Gaspelin et al. 2015; Coeperus & Quirk 2015), more or less explicitly setting out to test the emergence of adult-like mechanisms. Other studies, however, showing differences in the electrode or timing of EEG components between adults and children suggest that the two groups may instantiate attentional control through different neuro(cognitive) mechanisms. In this study, we were interested in the extent to which attentional control processes (TAC and MSE) that are present in adults, together with their brain mechanisms (here, modulations in the predominance of topographic maps, and thus recruited networks), are also present in children at different ages. What our current analyses could not reveal is what children’s spatially-selective brain mechanisms ‘look like’ at different ages, with child groups, and not adults, acting as a reference point. In the pursuit of understanding of the development of attentional control in real-world environments, the two perspectives complement each other, creating a more complete picture on attentional control development. For this reason, we have recently carried out a child-centric EN analysis of the developing visual and multisensory attentional control (Turoman et al., 2020b). In short, we found that when using children as a reference point, already 5-year-olds show evidence of visual top-down attentional control, as measured by TAC, albeit not via adult-like brain networks.

### 4. Conclusion

Taken together, our study revealed the developmental trajectory of a frequently studied visual attentional control mechanism that is task-set contingent attentional capture (TAC). We showed, both behaviourally and using an EN analytical framework, that TAC develops early in childhood (after the age of 5 years), and reaches adult-like state at 7 years of age. Though MSE, present in adults, was undetected in children’s behaviour or traditionally analysed EEG signals, an EN framework revealed spatially-selective brain mechanisms sensitive to the multisensory nature of distractors, at 8-9 years. Our findings underline the utility of combining traditional behavioural and EEG/ERP markers of visual attentional control with multivariate EEG analytical techniques for investigating the development of attentional control, and for identifying developmental differences and similarities in attentional control between adults and children.

## Acknowledgements

Acknowledgments

We thank The EEG Brain Mapping Core of the Center for Biomedical Imaging (CIBM) for providing the infrastructure. We would also like to thank Micah Murray for helpful comments during the conceptualisation of the study. We would also like to thank Noémie Kirscher for assistance with child IQ data collection and Louise Vasa for assistance with child EEG data analysis. Funding: This work was supported by the Pierre Mercier Foundation and the Swiss National Science Foundation (grant PZ00P1_174150) grants to P.J.M., as well as by the Fondation Asile des Aveugles.

## References

Astle, D. E., & Scerif, G. (2011). Interactions between attention and visual short-term memory (VSTM): What can be learnt from individual and developmental differences?. Neuropsychologia, 49(6), 1435–1445.

Bahrick, L. E., & Lickliter, R. (2000). Intersensory redundancy guides attentional selectivity and perceptual learning in infancy. Developmental psychology, 36(2), 190–201.

Barutchu, A., Crewther, D. P., & Crewther, S. G. (2009). The race that precedes coactivation: development of multisensory facilitation in children. Developmental science, 12(3), 464–473.

Broadbent, H. J., Osborne, T., Mareschal, D., & Kirkham, N. Z. (2019). Withstanding the test of time: Multisensory cues improve the delayed retention of incidental learning. Developmental science, 22(1), e12726.

Broadbent, H. J., White, H., Mareschal, D., & Kirkham, N. Z. (2018). Incidental learning in a multisensory environment across childhood. Developmental science, 21(2), e12554.

Brod, G., Bunge, S. A., & Shing, Y. L. (2017). Does one year of schooling improve children’s cognitive control and alter associated brain activation?. Psychological science, 28(7), 967–978.

Brunet, D., Murray, M. M., & Michel, C. M. (2011). Spatiotemporal analysis of multichannel EEG: CARTOOL. Computational intelligence and neuroscience, 2011, 2.

Buchs, G., Heimler, B., & Amedi, A. (2019). The Effect of Irrelevant Environmental Noise on the Performance of Visual-to-Auditory Sensory Substitution Devices Used by Blind Adults. Multisensory research, 32(2), 87–109.

Bunge, S. A., Dudukovic, N. M., Thomason, M. E., Vaidya, C. J., & Gabrieli, J. D. (2002). Immature frontal lobe contributions to cognitive control in children: evidence from fMRI. Neuron, 33(2), 301–311.

Cappe, C., Thut, G., Romei, V., & Murray, M. M. (2010). Auditory–visual multisensory interactions in humans: timing, topography, directionality, and sources. Journal of Neuroscience, 30(38), 12572–12580.

Casey, B. J., Tottenham, N., Liston, C., & Durston, S. (2005). Imaging the developing brain: what have we learned about cognitive development?. Trends in cognitive sciences, 9(3), 104–110.

Conférence intercantonale de l’instruction publique de la Suisse romande et du Tessin (CIIP). Plan d’études romand (PER), Aperçu des contenus (June 2012). Available at: https://www.plandetudes.ch/documents/10136/19192/cycle_1_webCIIP.pdf

Couperus, J. W., & Quirk, C. (2015). Visual search and the N2pc in children. Attention, Perception, & Psychophysics, 77(3), 768–776.

De Meo, R., Murray, M. M., Clarke, S., & Matusz, P. J. (2015). Top-down control and early multisensory processes: chicken vs. egg. Frontiers in integrative neuroscience, 9, 17.

Denervaud, S., Gentaz, E., Matusz, P. J., & Murray, M. M. (2020). Multisensory gains in simple detection predict global cognition in schoolchildren. Scientific Reports, 10, 1–11.

Donnelly, N., Cave, K., Greenway, R., Hadwin, J. A., Stevenson, J., & Sonuga-Barke, E. (2007). Visual search in children and adults: Top-down and bottom-up mechanisms. Quarterly Journal of Experimental Psychology, 60(1), 120–136.

Eimer, M. (1996). The N2pc component as an indicator of attentional selectivity. Electroencephalography and clinical neurophysiology, 99(3), 225–234.

Eimer, M. (2014). The neural basis of attentional control in visual search. Trends in Cognitive Sciences, 18(10), 526–535.

Eimer, M., Kiss, M., Press, C., & Sauter, D. (2009). The roles of feature-specific task set and bottom-up salience in attentional capture: an ERP study. Journal of Experimental Psychology: Human Perception and Performance, 35(5), 1316.

Fisher, A. V., Godwin, K. E., & Seltman, H. (2014). Visual environment, attention allocation, and learning in young children: When too much of a good thing may be bad. Psychological science, 25(7), 1362–1370.

Folk, C. L., Remington, R. W., & Johnston, J. C. (1992). Involuntary covert orienting is contingent on attentional control settings. Journal of Experimental Psychology: Human perception and performance, 18(4), 1030.

Folk, C. L., & Remington, R. W. (1998). Selectivity in distraction by irrelevant featural singletons: evidence for two forms of attentional capture. Journal of Experimental Psychology: Human perception and performance, 24(3), 847–858.

Gaspelin, N., Margett-Jordan, T., & Ruthruff, E. (2015). Susceptible to distraction: Children lack top-down control over spatial attention capture. Psychonomic bulletin & review, 22(2), 461–468.

Giard, M. H., & Peronnet, F. (1999). Auditory-visual integration during multimodal object recognition in humans: a behavioral and electrophysiological study. Journal of cognitive neuroscience, 11(5), 473–490.

Gori, M., Del Viva, M., Sandini, G., & Burr, D. C. (2008). Young children do not integrate visual and haptic form information. Current Biology, 18(9), 694–698.

Gori, M., Sandini, G., & Burr, D. (2012). Development of visuo-auditory integration in space and time. Frontiers in integrative neuroscience, 6, 77.

Hommel, B., Li, K. Z., & Li, S. C. (2004). Visual search across the life span. Developmental psychology, 40(4), 545.

Hwang, K., Velanova, K., & Luna, B. (2010). Strengthening of top-down frontal cognitive control networks underlying the development of inhibitory control: a functional magnetic resonance imaging effective connectivity study. Journal of Neuroscience, 30(46), 15535–15545.

Kiss, M., Jolicœur, P., Dell’Acqua, R., & Eimer, M. (2008). Attentional capture by visual singletons is mediated by top-down task set: New evidence from the N2pc component. Psychophysiology, 45(6), 1013–1024.

Konrad, K., Neufang, S., Thiel, C. M., Specht, K., Hanisch, C., Fan, J., & Fink, G. R. (2005). Development of attentional networks: an fMRI study with children and adults. Neuroimage, 28(2), 429–439.

Lehmann, D., & Skrandies, W. (1980). Reference-free identification of components of checkerboard-evoked multichannel potential fields. Electroencephalography and clinical neurophysiology, 48(6), 609–621.

Lewkowicz, D. J. (2014). Early experience and multisensory perceptual narrowing. Developmental psychobiology, 56(2), 292–315.

Lewkowicz, D. J., & Turkewitz, G. (1980). Cross-modal equivalence in early infancy: Auditory–visual intensity matching. Developmental psychology, 16(6), 597.

Luck, S. J., & Hillyard, S. A. (1994). Spatial filtering during visual search: evidence from human electrophysiology. Journal of Experimental Psychology: Human Perception and Performance, 20(5), 1000.

Massonnié, J., Rogers, C. J., Mareschal, D., & Kirkham, N. Z. (2019). Is classroom noise always bad for children? The contribution of age and selective attention to creative performance in noise. Frontiers in psychology, 10, 381.

Matusz, P. J., & Eimer, M. (2011). Multisensory enhancement of attentional capture in visual search. Psychonomic Bulletin & Review, 18(5), 904.

Matusz, P. J., Broadbent, H., Ferrari, J., Forrest, B., Merkley, R., & Scerif, G. (2015). Multi-modal distraction: Insights from children’s limited attention. Cognition, 136, 156–165.

Matusz, P. J., Key, A. P., Gogliotti, S., Pearson, J., Auld, M. L., Murray, M. M., & Maitre, N. L. (2018). Somatosensory plasticity in pediatric cerebral palsy following constraint-induced movement therapy. Neural plasticity, 2018.

Matusz, P. J., Merkley, R., Faure, M., & Scerif, G. (2019a). Expert attention: Attentional allocation depends on the differential development of multisensory number representations. Cognition, 186, 171–177.

Matusz, P. J., Turoman, N., Tivadar, R. I., Retsa, C., & Murray, M. M. (2019b). Brain and cognitive mechanisms of top–down attentional control in a multisensory world: Benefits of electrical neuroimaging. Journal of cognitive neuroscience, 31(3), 412–430.

Maylor, E. A., & Lavie, N. (1998). The influence of perceptual load on age differences in selective attention. Psychology and aging, 13(4), 563.

Melinder, A., Gredeback, G., Westerlund, A., & Nelson, C. A. (2010). Brain activation during upright and inverted encoding of own-and other-age faces: ERP evidence for an own-age bias. Developmental Science, 13, 588–598.

Michel, C. M., & Koenig, T. (2018). EEG microstates as a tool for studying the temporal dynamics of whole-brain neuronal networks: a review. Neuroimage, 180, 577–593.

Murray, M. M., Brunet, D., & Michel, C. M. (2008). Topographic ERP analyses: a step-by-step tutorial review. Brain topography, 20(4), 249–264.

Murray, M. M., Thelen, A., Thut, G., Romei, V., Martuzzi, R., & Matusz, P. J. (2016). The multisensory function of the human primary visual cortex. Neuropsychologia, 83, 161–169.

Murray, M. M., Matusz, P. J., & Amedi, A. (2015). Neuroplasticity: unexpected consequences of early blindness. Current Biology, 25, R998–R1001.

Murray, M.M. and Wallace, M.T., eds (2012) The Neural Bases of Multisensory Processes, CRC Press

Neil, P. A., Chee-Ruiter, C., Scheier, C., Lewkowicz, D. J., & Shimojo, S. (2006). Development of multisensory spatial integration and perception in humans. Developmental science, 9(5), 454–464.

Oh-uchi, A., Kawahara, J. I., & Sugano, L. (2010). Attentional capture and metaattentional judgment: a study of young children, parents, and university students. Psychologia, 53(2), 114–124.

Petrini, K., Jones, P. R., Smith, L., & Nardini, M. (2015). Hearing where the eyes see: Children use an irrelevant visual cue when localizing sounds. Child development, 86(5), 1449–1457.

Perrin, F., Pernier, J., Bertnard, O., Giard, M. H., & Echallier, J. F. (1987). Mapping of scalp potentials by surface spline interpolation. Electroencephalography and clinical neurophysiology, 66(1), 75–81.

Rueda, M. R., Fan, J., McCandliss, B. D., Halparin, J. D., Gruber, D. B., Lercari, L. P., & Posner, M. I. (2004). Development of attentional networks in childhood. Neuropsychologia, 42(8), 1029–1040.

Santangelo, V., & Spence, C. (2007). Multisensory cues capture spatial attention regardless of perceptual load. Journal of Experimental Psychology: Human Perception and Performance, 33(6), 1311.

Sawaki, R., & Luck, S. J. (2010). Capture versus suppression of attention by salient singletons: Electrophysiological evidence for an automatic attend-to-me signal. Attention, Perception, & Psychophysics, 72(6), 1455–1470.

Scerif, G., Kotsoni, E., & Casey, B. J. (2006). Functional Neuroimaging of Early Cognitive Development. Handbook of Functional Neuroimaging of Cognition, 351.

Schwartze, M., Rothermich, K., Schmidt-Kassow, M., & Kotz, S. A. (2011). Temporal regularity effects on pre-attentive and attentive processing of deviance. Biological psychology, 87(1), 146–151.

Shimi, A., & Scerif, G. (2017). Towards an integrative model of visual short-term memory maintenance: Evidence from the effects of attentional control, load, decay, and their interactions in childhood. Cognition, 169, 61–83.

Shimi, A., Nobre, A. C., Astle, D., & Scerif, G. (2014a). Orienting attention within visual short-term memory: Development and mechanisms. Child development, 85(2), 578–592.

Shimi, A., Kuo, B. C., Astle, D. E., Nobre, A. C., & Scerif, G. (2014b). Age group and individual differences in attentional orienting dissociate neural mechanisms of encoding and maintenance in visual STM. Journal of Cognitive Neuroscience, 26(4), 864–877.

Shimi, A., Nobre, A. C., & Scerif, G. (2015). ERP markers of target selection discriminate children with high vs. low working memory capacity. Frontiers in systems neuroscience, 9, 153.

Stein, B. E., & Meredith, M. A. (1993). The merging of the senses. The MIT Press.

Sun, M., Wang, E., Huang, J., Zhao, C., Guo, J., Li, D., & Song, Y. (2018). Attentional selection and suppression in children and adults. Developmental science, 21(6), e12684.

Talsma, D., Senkowski, D., Soto-Faraco, S., & Woldorff, M. G. (2010). The multifaceted interplay between attention and multisensory integration. Trends in cognitive sciences, 14(9), 400–410.

Ten Oever, S., Romei, V., van Atteveldt, N., Soto-Faraco, S., Murray, M. M., & Matusz, P. J. (2016). The COGs (context, object, and goals) in multisensory processing. Experimental brain research, 234(5), 1307–1323.

Tivadar, R. I., & Murray, M. M. (2019). A primer on electroencephalography and event-related potentials for organizational neuroscience. Organizational Research Methods, 22(1), 69–94.

Trick, L. M., & Enns, J. T. (1998). Lifespan changes in attention: The visual search task. Cognitive Development, 13(3), 369–386.

Tsujimoto, S. (2008). The prefrontal cortex: Functional neural development during early childhood. The Neuroscientist, 14(4), 345–358.

Turoman, N., Tivadar, R., Retsa, C., Murray, M. M., & Matusz, P. J. (2020a). How we pay attention in naturalistic settings. BioRxiv. DOI:10.1101/2020.07.30.229617

Turoman, N., Tivadar, R. I., Retsa, C., Maillard, A. M., Scerif, G., & Matusz, P. J. (2020b). Uncovering the mechanisms of real-world attentional control over the course of primary education. BioRxiv. DOI:10.1101/2020.10.20.342758

Van der Burg, E., Talsma, D., Olivers, C. N., Hickey, C., & Theeuwes, J. (2011). Early multisensory interactions affect the competition among multiple visual objects. Neuroimage, 55(3), 1208–1218.

Wechsler, D. (2012). Wechsler preschool and primary scale of intelligence (4th ed.). Bloomington, MN: Pearson.

Wechsler, D. (2014). Wechsler intelligence scale for children-fifth edition. Bloomington, MN: Pearson.

Yang, Q., Bucci, M. P., & Kapoula, Z. (2002). The latency of saccades, vergence, and combined eye movements in children and in adults. Investigative Ophthalmology & Visual Science, 43(9), 2939–2949.

