## Supplemental Materials for "The development of attentional control mechanisms in multisensory environments"

***Supplementary Methods & Results***

Turoman, Tivadar, Retsa, Maillard, Scerif, Matusz

**Methods**

**EEG preprocessing**

**EEG data interpolation**

Average numbers of epochs removed were as follows: 11% in adults, 11% in 9-year olds, 8% in 7-year olds, and 14% in 5-year olds. Average numbers of electrodes interpolated per participant were as follows: 8 in adults (6% of the total electrode montage), 11 in 9-year olds (8% of the total electrode montage), 10 in 7-year olds (8% of the total electrode montage), and 12 in 5-year olds (10% of the total electrode montage).

**Results**

**Supplemental behavioural results**

**Behavioural results: RTs corrected for age-related slowing**

After correcting for cognitive slowing, the ANOVA revealed no significant differences in mean RT by Age, *F* = 0.01, *p* = 0.99, which is not unexpected, given that we scaled RT data to take into account differential overall RT for different participants’ ages. As in the corrected RT data analysis, Age did not interact with any other factors (all *F's* < 2, *p*’s > 0.1).

In 9-year-olds, behavioural capture effects were reliable, with responses faster on trials where the cue and target location were the same (839ms) versus when they were different (870ms), *F*_(1, 25)_ = 65.6, *p* < 0.001, η_p_² = 0.7. Like in adults, behavioural capture effects were modulated by Cue Colour, *F*_(1, 25)_ = 20.3, *p* < 0.001, η_p_² = 0.5. This effect was driven by significant cueing effects for the TCC distractors (56ms, *t*_(25)_ = 8.4, *p* < 0.001), but not the NCC distractors (6ms, *t*_(25)_ = 1, *p* = 0.6). As in the case of raw RTs, there was no evidence for modulation of capture effects by Cue Modality, *F*_(1, 25)_ = 1.4, *p* = 0.3.

Similarly, in 7-year-olds, behavioural capture effects were reliable, with faster responses on trials where the cue and target location were the same (1109ms) versus when they were different (1114ms), *F*_(1, 37)_ = 11, *p* < 0.001, η_p_² = 0.2. These capture effects were also modulated by Cue Colour, *F*_(1, 37)_ = 8, *p* = 0.008, η_p_² = 0.2, such that capture effects were significant for the TCC distractors (55ms, *t*_(37)_ = 4.3, *p* < 0.001), but not the NCC distractors (7ms, *t*_(37)_ = 0.03, *p* = 1). Again, capture effects were not modulated by Cue Modality, *F*_(1, 37)_ = 2.1, *p* = 0.2.

In 5-year-olds, behavioural capture effects were reliable, with faster RTs on trials where the cue and target location were the same (1312ms) versus when they were different (1343ms), *F*_(1, 27)_ = 4.4, *p* = 0.045, η_p_² = 0.1. As in the case for raw RTs, behavioural capture was not modulated by Cue Colour (*F*_(1, 27)_ = 1, *p* = 0.3), or by Cue Modality (*F*_(1, 27)_ = 0.03, *p* = 0.9).

**Supplemental N2pc results**

**Additional canonical analysis results**

Across the child groups, there were no significant main effects and interactions (*F*’s < 1), except for the main effect of Cue Modality. In 9-year-olds, the main effect of Cue Modality, *F*_(1, 25)_ = 60.5, *p* < 0.001, η_p_² = 0.7, was driven by more positive ERP amplitudes for AV distractors (1.9μV) than for V distractors (-0.3μV). Likewise, in 7-year-olds, the main effect of Cue Modality, *F*_(1, 37)_ = 35, *p* < 0.001, η_p_² = 0.5, was driven by more positive ERP amplitudes for AV distractors (0.9μV) than for V distractors (-1.6μV). Meanwhile, in 5-year-olds, the main effect of Cue Modality had the level of a nonsignificant trend, *F*_(1, 27)_ = 3.6, *p* = 0.07, η_p_² = 0.1, although, even here, numerically, ERP amplitudes were more positive for AV distractors (2.2μV) than for V distractors (0.3μV).

**Data-driven approach**

Here, N2pc time-window and electrode choices were based on adult EEG data, rather than taken from the literature, and were obtained from the following series of steps. We first computed a contralateral-ipsilateral difference waveform by subtracting the voltage amplitudes over the contralateral hemifield (left side of the scalp) from voltage amplitudes over the ipsilateral hemifield (right side of the scalp) for the experimental condition which best resembled the stimulus settings in which the N2pc is traditionally observed, i.e., the target matching visual condition - TCCV. Individual difference waveforms for this condition were grand averaged. To obtain an N2pc time-window, in the grand-averaged TCCV difference waveform, we identified the post-cue time-period at which the DISS measure was stable, suggesting a single set of active brain networks that would underlie the N2pc. The resulting time window was 154-300ms (147ms duration). Next, to identify the appropriate electrode sites, in the grand-averaged TCCV ERP, we identified the locations on the scalp with the highest negative voltage amplitudes over the above N2pc time-windows. The resulting electrodes were e59/e91 (contralateral/ipsilateral, respectively). We then extracted mean voltage amplitudes from the above electrodes in the above time-window for each of the four age groups, and submitted the data to separate 3-way repeated-measures ANOVAs, with within-subject factors: Cue Colour (TCC vs. NCC), Cue Modality (V vs. AV), and Contralaterality (Contralateral vs. Ipsilateral).

In adults, there was a reliable N2pc, supported by a main effect of Contralaterality, *F*_(1, 38)_ = 17.6, *p* < 0.001, η_p_² = 0.3. The mean of the overall N2pc amplitude was -0.5μV, with the contralateral and ipsilateral mean amplitude being -0.6μV and -0.1μV, respectively. There was a 2-way Contralaterality x Cue Colour interaction, *F*_(1, 38)_ = 10.7, *p* = 0.002, η_p_² = 0.2, such that N2pc’s were reliable for TCC distractors (-0.7μV), but only at the level of a nonsignificant trend for NCC distractors (-0.2μV). The 2-way Contralaterality x Cue Modality interaction was not significant (*F* < 1). Notably, with the parameters for canonical N2pc analyses chosen in a more data-driven fashion, the 3-way Contralaterality x Cue Colour x Cue Modality interaction was no longer significant, *F*_(1, 38)_ = 0.03, *p* = 0.9, η_p_² = 0.001. There were significant main effects of Cue Colour and Cue Modality. The main effect of Cue Colour, *F* = 10.7, *p* = 0.002, η_p_² = 0.2, was driven by larger ERP amplitudes for TCC distractors (-0.5μV) than for NCC distractors (-0.2μV). Meanwhile the main effect of Cue Modality, *F* = 6.3, *p* = 0.016, η_p_² = 0.1, was driven by larger ERP amplitudes for V distractors (-0.5μV) than for AV distractors (-0.2μV).

In 9-year-olds, only the main effect of Cue Modality was significant, *F*_(1, 25)_ = 45.8, *p* < 0.001, η_p_² = 0.7, and was driven by larger ERP amplitudes for V distractors (-0.1μV) than for AV distractors (1.7μV). Importantly, the main effect of Contralaterality did not reach statistical significance, *F*_(1, 25)_ = 0.9, *p* = 0.4, and neither did the other main effects or interactions (all *F’s* < 1).

Similarly, in 7-year-olds, only the main effect of Cue Modality was significant, *F*_(1, 37)_ = 42.4, *p* < 0.001, η_p_² = 0.5, and driven by larger ERP amplitudes for V distractors (-1.3μV) than for AV distractors (0.9μV). Meanwhile the main effect of Contralaterality, *F*_(1, 37)_ = 0.1, *p* = 0.7, and other main effects or interactions were not significant (all *F’s* < 1).

In 5-year-olds, the main effect of Cue Modality was now significant, *F*_(1, 27)_ = 9.4, *p* = 0.05, η_p_² = 0.3, and was, like in the older age groups, driven by larger ERP amplitudes for V distractors (-0.2μV) than for AV distractors (1.6μV). Meanwhile the main effect of Contralaterality, *F*_(1, 27)_ = 0.3, *p* = 0.6, and other main effects or interactions were not (all *F’s* < 1).

**Supplemental GFP results**

In adults, the main effect of Cue Colour, *F*_(1, 38)_ = 2.8, *p* = 0.1, *ηp²* = 0.07, and of Cue Modality, *F*_(1, 38)_ =0.1, *p* = 0.7, *ηp²* = 0.003 did not reach statistical significance, and neither did the two-way interaction between these factors, *F*_(1, 38)_ = 0.5, *p* = 0.5, *ηp²* = 0.01. Similarly, in 9-year-olds, the main effect of Cue Colour, *F*_(1, 25)_ = 0.1, *p* = 0.7, *ηp²* = 0.004, and of Cue Modality, *F*_(1, 25)_ = 0.8, *p* = 0.4, *ηp²* = 0.03, as well as the two-way interaction between these factors, *F*_(1, 25)_ = 0.05, *p* = 0.8, *ηp²* = 0.002, were not significant. In 7-year-olds, the main effect of Cue Colour reached a nonsignificant trend**,** *F*_(1, 37)_ = 3.9 *p* = 0.06, *ηp²* = 0.1. However, the main effect of Cue Modality, *F*_(1, 37)_ = 1.5, *p* = 0.2, *ηp²* = 0.004, and the two-way interaction between Cue Colour and Cue Modality were not significant, *F*_(1, 37)_ = 6, *p* = 0.02, *ηp²* = 0.1. Finally, in 5-year-olds, the main effect of Cue Colour, *F*_(1, 27)_ = 1.2, *p* = 0.3, *ηp²* = 0.04, and of Cue Modality, *F*_(1, 27)_ = 0.3, *p* = 0.6, *ηp²* = 0.01, as well as the two-way interaction between these factors, *F*_(1, 27)_ = 0.1, *p* = 0.7, *ηp²* = 0.005, were not significant.

**Supplemental figures**

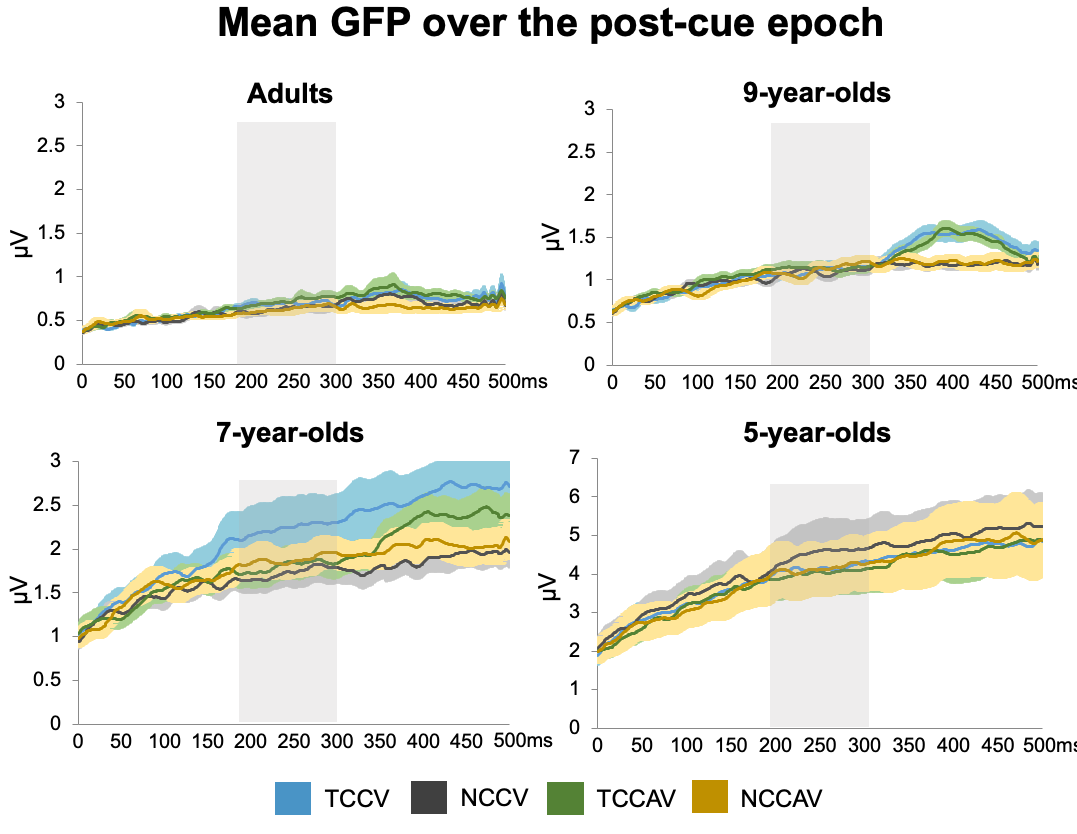

**Supplementary Figure 1.** Overall GFP for each of the 4 cue conditions (represented according to the figure legend) per age-group, plotted across the entire post-cue time-window. Thick lines represent mean GFP’s while the surrounding lighter coloured fields represent the standard error of the mean. For reference, the boundaries of the canonical N2pc time-window are highlighted in grey. As visible from overlapping means and error areas, there was no significant main effects and interactions in any of the tested age groups.
